# Distinct movement patterns generate stages of spider web-building

**DOI:** 10.1101/2021.05.24.444987

**Authors:** Abel Corver, Nicholas Wilkerson, Jeremiah Miller, Andrew Gordus

## Abstract

The geometric complexity and stereotypy of spider webs have long generated interest in their algorithmic origin. Like other examples of animal architecture, web construction is the result of several assembly phases that are driven by distinct behavioral stages coordinated to build a successful structure. Manual observations have revealed a range of sensory cues and movement patterns used during web construction, but methods to systematically quantify the dynamics of these sensorimotor patterns are lacking. Here, we apply an analytical pipeline to quantify web-making behavior of the orb-weaver *Uloborus diversus*. Position tracking revealed stereotyped stages of construction that could occur in typical or atypical progressions across individuals. Using an unsupervised clustering approach, we identified general and stage-specific leg movements. A Hierarchical Hidden Markov Model revealed that stages of web-building are characterized by stereotyped sequences of actions largely shared across individuals, regardless of whether these stages progress in a typical or atypical fashion. Web stages could be predicted based on action-sequences alone, revealing that web-stages are a physical manifestation of underlying behavioral phases.

**Highlights:**

1. Spider centroid trajectories indicate stereotyped progression of web-building stages.
2. Unsupervised movement clustering reveals a shared set of movements which correspond to previously defined behaviors that define web-making across individuals.
3. Stages of web-building use both stage-specific and non-specific behaviors.
4. Stereotyped and distinct action sequences are predictive of stages of web-building.

## Introduction

In Ovid’s *Metamorphoses*, orb-weavers are the descendants of Arachne, a human weaver condemned to the form of a spider for weaving tapestries more beautiful than the Gods’^1^. In the pantheon of animals, the allure of this impressive behavior, both scientifically and aesthetically, lies in the completed product – the web – which is a physical record of many aspects of a spider’s behavior^2^. Internal states such as satiety, sexual arousal, or aggression strongly influence behavior^3^, however a challenge for many behavioral paradigms is that these states must be inferred indirectly, often from stochastic behavioral state-switching^3^. Unlike web-building, these behaviors do not provide a record of the transitions between behavioral states. For example, foraging is composed of dwelling and roaming states which are sampled randomly^4^. These states can be inferred probabilistically by the likelihoods of behavioral metrics such as velocity and turning rates, but these observable metrics themselves do not directly reflect the internal state of the animal. Instead, if the internal state is hunger, the animal is *more likely to* roam in search of food, and if it is sated it is *more likely* to dwell with low velocity. However, these boundaries are not absolute; hungry animals also dwell, and sated animals also roam, but the probability of these behaviors are different depending on the internal state.

Ideally, a paradigm where the animal provided a record of their intent would simplify state identification, making more precise subsequent biological assays to understand how these states are encoded neuronally. An advantage of quantifying orb-weaving behavior is that orb web construction occurs in behavioral phases that are easily defined by the spider’s trajectory and web geometry^5^. Despite a diversity of species and preferred environments, all orb-weavers share the same progression of web-construction phases that are the result of many shared behaviors such as walking and silk-extrusion. Though some of these behaviors occur in all stages of web-building^6^, how these behaviors are coordinated differs across web-construction phases^7^. In principle, different phases of web-building should be predictable based on these different behavioral strategies, independent of the spider’s trajectory. While the general phases of assembly have been well-documented^8^, a detailed quantification of the behaviors that embody these phases, and their flexibility, has not been systematically addressed^9^. Even though the sequence of web-construction phases is fairly stereotyped, several instances of atypical webs have been observed^10^. Such atypical webs include so-called “senile webs”, but the changes in behavior that lead to these atypical webs is not well understood^11^.

Here, we develop an analytical framework for defining web-building behaviors and identify stereotyped action sequences that characterize the construction of a spider’s orb-web. While the typical progression of web stages was observed, several atypical progressions were also noticed. This analysis revealed quantitatively that the different phases of web-building are plastic. Even though web-construction often follows a typical sequence of construction phases, atypical progression can occur even in the context of apparently complete coverage of the prior construction phase. Each phase of web-building can be explicitly defined by shared and unique underlying behaviors, and these definitions are the same for typical and atypical web-constructions. Thus, different phases of web-building are an external record of different underlying behavioral states.

## Results

### The spider’s trajectory reflects each stage of web-building

While most orb-weaving spiders are members of the family Araneidae^12,13^, the Uloboridae also produce orb-webs with the same underlying structural motifs^14^. Most orb-weavers are primarily active during the spring and summer, but other species like *Uloborus diversus*^15^ are active throughout the year^6^. Due to this prolonged activity, as well as other features like their small size and tolerance of conspecifics, we chose to use *U. diversus* for our studies (Figure 1A). While *U. diversus* can build webs within a variety of geometries, it is a nocturnal animal that prefers to build horizontal webs in complete darkness, so we built a behavior arena to accommodate these preferences (Figures 1B-D).

**Figure 1:**
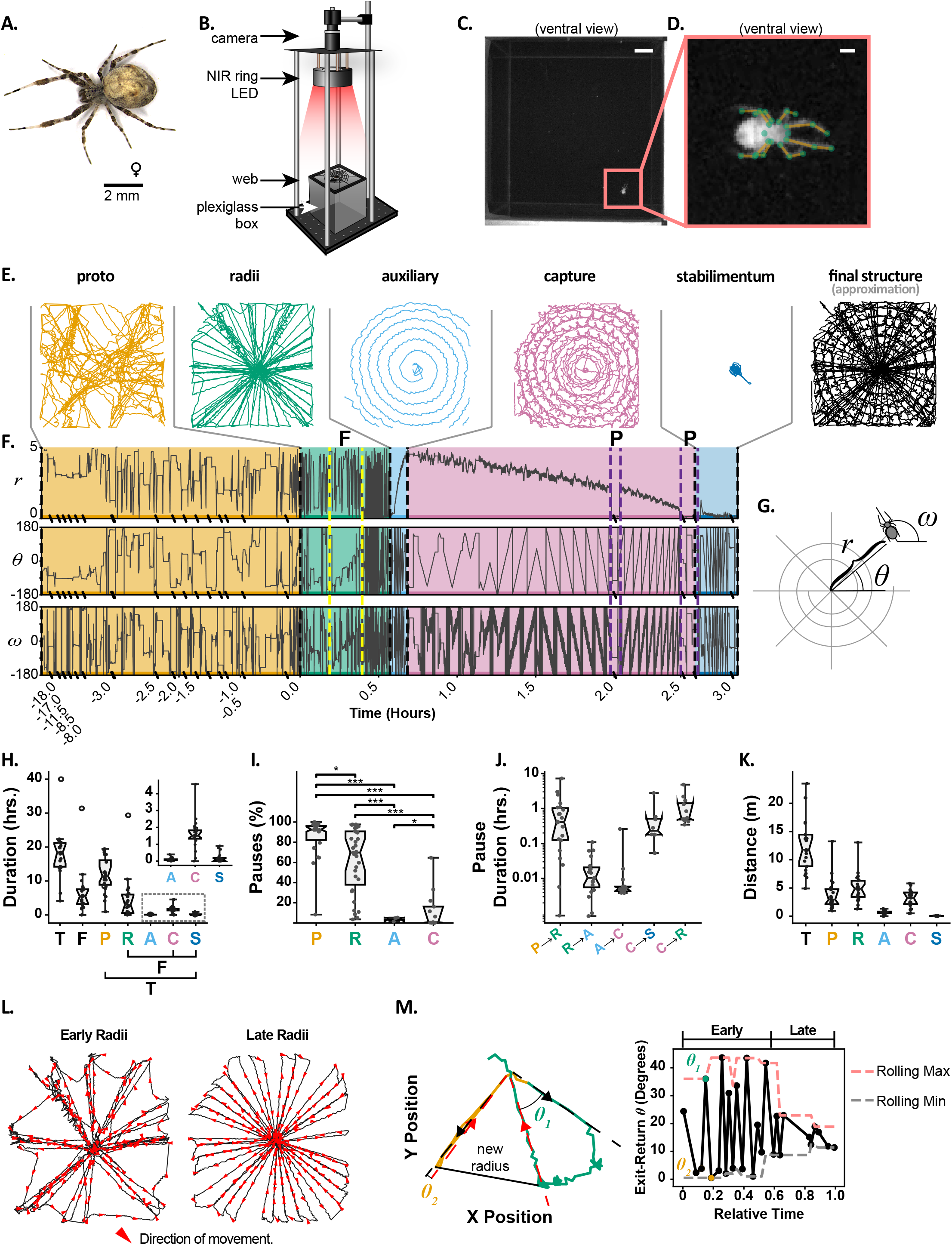
Centroid trajectories indicate stereotyped progression of web-building stages. A. Representative adult female used in this study. B Experimental setup: Spiders assembled webs on a plexiglass box under NIR (880 nm) illumination and were recorded by a USB CMOS camera at 50 Hz. C. Example of typical frame acquired. Scale bar = 1 cm. D. Example of centered and rotated frame, with limb annotation. Scale bar = 1 mm. Green dots indicate anatomical points that were tracked in later analyses. E. Example centroid trajectories from different web stages for a single web. The final structure approximation is based on superimposing the radii, capture and stabilimentum stages. F. Radial position and orientation coordinates for the web progression in (E). Colors correspond to the stages of web-building in (E). “P” indicates pauses during the capture spiral, and “F” indicates frame construction. Hatch marks along the time-axis indicate time compressions where the spider paused for extended durations. G. Polar coordinate reference for coordinate time series in (F). H. Stage durations for all recordings: Proto-web (P), Radii (R), Auxiliary Spiral (A), Capture Spiral (C) and Stabilimentum (S). Colors correspond to the stages of web-building in (E). The total duration (T) is obtained by adding all stage durations, and the final-structure duration (F) is obtained by adding only the radii, capture spiral and stabilimentum durations. Open circles correspond to an outlier recording in which the spider completed two complete auxiliary and capture spirals. The first 10% of distance traveled in the proto-web was not included in the proto-web duration. Notches represent the 95% confidence interval of the median. The inset displays the same spiral and stabilimentum durations on a smaller axis. I. Pause durations as a percentage of stage duration, computed over the [10%, 90%] interval of distance traveled in each stage to exclude pauses occurring during stage transitions. Notches represent the 95% confidence interval of the median. P-values based on Bonferroni-corrected two-sided Mann-Whitney rank tests (*: p < 0.05, **: p < 0.01, ***: p < 0.001). J. Total pause durations occurring between stages. The stage transition interval was defined as starting after 90% of distance traveled in the preceding stage and ending at 10% of distance traveled in the subsequent stage. Capture spiral to Radii (C→R) transitions represent atypical transitions as shown in Figure 2. Notches represent the 95% confidence interval of the median. K. Total distance traveled by stage and the total (T) distance traveled for each recording. Notches represent the 95% confidence interval of the median. L. Radii stage for an example recording, demonstrating an early and late phase. Red arrows indicate direction of travel. M. Left: Two consecutive radial exit-return trajectories, the first (green) spanning a large angle (⊝_1_ ≫ 0) followed by a second exit-return trajectory (yellow) re-traversing the same line (⊝_2_ ≈ 0). Black and red arrows indicate exit and return trajectories, respectively. Right: Sequence of exit-return angles over the course of the radii stage, for the same example recording. Yellow and green markers correspond to same-colored trajectories on the left. Dashed red and black lines represent the rolling (window=5) maximum and minimum, respectively.

The spider’s behavior was recorded for an average of 24 hours, spanning the entirety of web-constructions which lasts for several hours, at a framerate of 50 Hz to allow tracking of fast leg movements. Tracking the spider’s position alone revealed the common stereotypic trajectory most orb-weavers take when building an orb-web and has been a common approach used to track the spider’s progression through different orb-web phases^5^ (Figure 1E-H). Since the spider is constantly producing silk during web-construction, the position of the spider serves as a useful – though imperfect^5^ – proxy for silk location. As with most orb-weavers, the typical progression of orb-weaving with *U. diversus* starts with a proto-web, followed by the construction of the radii and frame, then an auxiliary spiral followed by a capture spiral. For some species such as *U. diversus*, an optional stabilimentum is also frequently observed^16^(Figures 1E-F, Supplementary Movie 1).

Like many orb-weavers, *U. diversus* spends a considerable amount of time exploring the arena and first builds a disorganized web called the “proto-web” (orange trajectories in Figures 1E,F). These webs have no obvious regularity, are mostly not part of the final structure, and are built with long irregular pauses that can last as long as 8 hours in our recordings (Figures 1H-K). It is thought that this stage of web-building is an exploratory phase where the spider assesses the structural integrity of its surroundings and locates anchor points for the final web^17^. This stage often ended with a prolonged pause that could last from minutes to hours (Figure 1J).

Once the spider progressed to radii and frame construction, it initially removed most of the proto-web, simultaneously adjusted some of the proto-web lines to serve as radii, and built the frame (Figure 1L). This early phase of radii construction often involved outward walks along a radius line and return along a newly anchored line (green trajectory in Figure 1M). This is characteristic of building new radii by anchoring silk at the hub, walking outward along a prior radius, then anchoring the new silk on the frame at an angular distance from the prior line and returning to the hub along this new line. The construction of the frame is often the result of anchoring silk on the periphery prior to the return, and then walking outward along a prior radius, anchoring the frame silk at the end of the line, and then returning to the hub^6^ (Figure 1M, orange trajectory). This early phase of radii construction could be defined by calculating the angle of exit and return from the hub (Figure 1M), as well as the gradual change in the spider’s angular position (Figure 1F, yellow outline in radii stage). We observed a tendency of the spider early in radii construction to alternate exit-return trajectories along the same line with exit-return trajectories that spanned a large angle (Figures 1L-M). The latter half of radii construction was characterized by a smaller and more consistent angular exit-return span as the spider progressed to a more regular construction of radii (Figures 1L-M).

Once the radii and frame were assembled, the spider would pause briefly, then spiral outward from the hub to create the auxiliary spiral (cyan trajectories in Figures 1E,F). This stage only lasted a few minutes with few pauses (Figures 1H-J). This stage is thought to stabilize the web structure for the subsequent construction of the capture spiral. The auxiliary spiral is temporary and was taken down as the spider spiraled inward from the periphery to create the capture spiral (pink trajectories in Figures 1E,F)., In some instances, a feature called a stabilimentum was added (blue trajectories in Figures 1E,F), which is thought to either camouflage the spider, attract prey, or visually deter collision by birds^18^. Once the spider finished the stabilimentum, it remained at the hub, sometimes for several days, as it awaited prey capture.

While centroid tracking easily discriminated transitions between most stages, the transition between proto-web and radii was more gradual and less distinct. This does not appear to be a limitation of centroid tracking, but rather reflects the inherently non-discrete boundary between these stages. During the initial phase of proto-web construction, nearly all the silk that is laid down is eventually removed. However, as this stage progresses, radii that will be part of the final structure are built or modified while the spider is still removing the proto-web. During this early stage of radii construction, the spider frequently traverses radii multiple times as it adjusts the web. Only when the proto-web has been completely removed and the frame assembled does the spider focus solely on radii construction and engage in the highly regular assembly of additional radii (Figure 1L).

All orb-weavers follow this progression of web stages^19^. While the stereotypy of this behavior has been well-studied, spiders do not necessarily rigidly transition through these stages, and will often interrupt a step to explore or modify the web, and then continue building (Figure 2A, Supplementary Figure 1A)^10^. Of the 21 webs recorded, the capture spiral was interrupted by a radial exploration stage twice, and additional radii stages followed the capture spiral or stabilimentum 9 times. This may be a type of error-assessment, since web damage will also disrupt web-building progression, followed by an attempt to fix the damage, and then a return to the prior web-building stage^6,20^. However, no obvious damage was observed in these experiments, though features like altered tension were not observable in this assay. These additional radii stages did not appear to be triggered by incomplete construction of prior radii as the density of radii at the beginning of these atypical stages were no different than typical radii density (Figure 2B). Most or all digressions from capture spiral construction appeared to be exploratory, since the radial trajectories of the spiders followed prior trajectories, indicating they were walking along previously built lines (Figure 2C). Alternatively, these additional radii stages may have been periods of “break-and-reel” when a spider will replace a prior radius with a new one by anchoring silk at the hub and walking along the old radius, and then breaking the old line and reeling in the new silk to anchor it to the frame^14^. In one unusual recording, the spider completed a secondary auxiliary and capture spiral stage, enlarging its prior completed web structure (Supplementary Figure 1A, Spider A4; Figure 6F).

**Figure 2:**
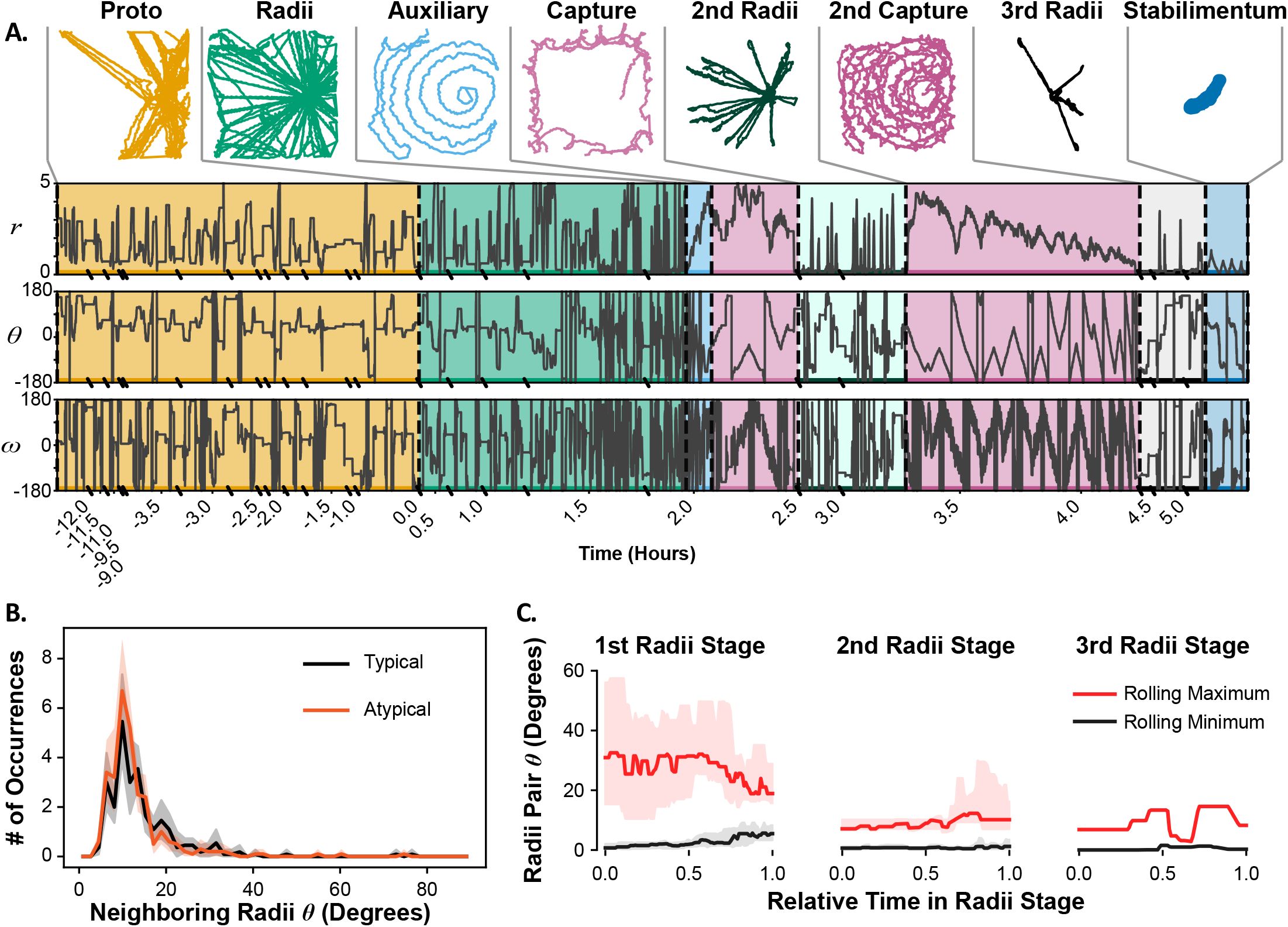
Atypical web stage progression. A. Top: Centroid trajectories for an atypical stage progression. Bottom: Radial position and angular coordinates for the stage progression shown above, with corresponding colors. The polar coordinate reference is as in Figure 1.G. B. Histogram of neighboring radii spacing after completion of the first radii stage. Histogram bins were based on the maximum of the Freedman-Diaconis rule applied to typical and atypical recordings separately. Shaded ribbons represent the 95% confidence interval of the mean for each bin. C. Rolling (window=5) minimum (black line) and maximum (red line) of the angular span of exit-return radial trajectory segments, as in Figure 1.M, averaged across recordings, split out by 1^st^ radii stage (the typical progression) and subsequent repetitions of this stage occurring in atypical recordings. Shaded regions correspond to the [25, 75] percentile confidence interval. Only one web had a 3^rd^ radii stage.

Atypical progressions may be due to atypical cues that trigger each stage of web-building. Even under constant conditions, an individual spider will build similar, but not identical, webs over several days (Supplementary Figure 1A). Tracking the position of the spider captures these differences, but it does not reveal the underlying differences in behavior that lead to these changes. One possibility is that atypical stage progression is the result of atypical underlying movement patterns. Alternatively, the underlying movements may be shared between typical and atypical progressions, and the atypical progressions are due to alternative internal or external sensory cues. To discriminate between these two possibilities, a more detailed understanding of the spider’s behavior is necessary.

### Spiders use a common repertoire of behaviors in web-building

To investigate spider behavior at a finer spatiotemporal scale, we tracked 26 points on its body: the base, femur and tibia of every leg, as well as the anterior- and posterior-most points of the prosoma (Figure 3A). To do this, 10,000 frames were manually annotated and used to entrain two limb-tracking convolutional neural networks (CNNs), LEAP^21^ and DeepLabCut^22^. Both CNNs performed similarly (8.2 and 7.6 mean pixel error for LEAP and DeepLabCut respectively, 4.5% and 3.3% errors ≥25 pixels respectively), and DeepLabCut was chosen for all subsequent analysis (Supplementary Figure 3A). Many of the largest errors occurred for medial leg coordinates. Due to this variance, and the more frequent use of anterior and posterior legs for web exploration and manipulation^6^, medial legs were ignored for subsequent analyses.

**Figure 3:**
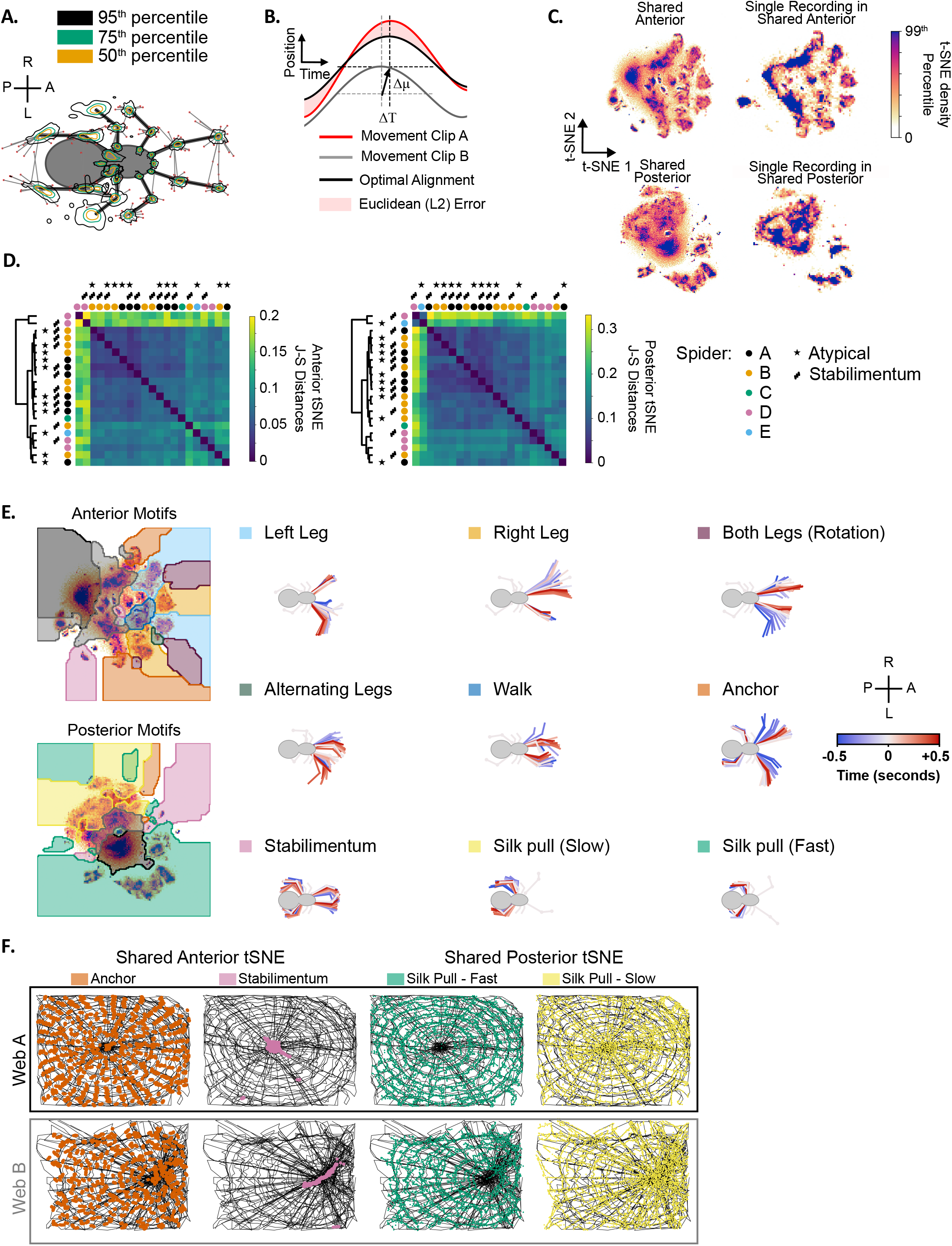
Unsupervised movement clustering reveals a shared set of movement motifs that characterize web-making across individuals. A. Confidence intervals of DeepLabCut-predicted limb coordinates. Tracking error vectors are superimposed onto an arbitrary reference posture. 95th (black), 75th (green) and 50th (yellow) percentile contours are displayed. Errors outside the 95th percentile are displayed as red dots. Anatomical axes: Anterior (A), Posterior (P), Right (R), Left (L). B. Euclidean error used for t-SNE distance metric. Before computing the Euclidean distance between two movement clips, every coordinate time series was mean-centered (dashed gray to dashed black horizontal lines, Δμ) and shifted in time (dashed gray to dashed black vertical lines, ΔT). C. Shared t-SNE embeddings averaged across recordings (Left) versus an example individual t-SNE embedding in the shared embedding space (Right). Densities for individual recordings were clipped to their 99^th^ percentile to show structure before averaging across recordings. Clipping prevents highly sampled (e.g. stationary) states from dominating the displayed embedding. D. Similarity of anterior and posterior embeddings across recordings, based on the Jensen-Shannon (J-S) distances. Dendograms based on hierarhical clustering of J-S distances. Circles colored according to individual spiders. Markers indicate atypical recordings and those with a stabilimentum. E. Left: Locations of movement motifs in t-SNE space for anterior and posterior limb trajectories. Right: Ventral view of representative 1-second limb trajectories corresponding to movement clusters on the left. Movements that were detected by both anterior and posterior limb embeddings are superimposed onto the same skeleton for illustration only. Anatomical axes: Anterior (A), Posterior (P), Right (R), Left (L). F. Spatial occurrences of 4 example behavioral motifs for two web recordings. Anchoring and Stabilimentum locations are based on anterior limb movements. Slow and fast silk pulling was signaled by posterior limb movements.

Since behavior is not simply static limb postures, but the movement of these postures in time, we initially attempted to capture this movement by passing a wavelet transform over the coordinates, as done previously for other invertebrates and vertebrates^23,24^. This approach defines each point in time as a vector of movement frequencies for each tracked coordinate, resulting in a multi-dimensional space of possible limb frequencies. However, we found the often irregular, and less sinusoidal, spider behaviors produced noisy wavelet profiles, unlike repetitive motions, like grooming or walking in flies and mice in a featureless environment, which produce a reliable signal in Fourier space (Supplementary Figure 3B). As an alternative approach, we chose to define each frame by an 880-ms interval centered on that frame (Figure 3B). While this fundamentally limited our temporal resolution of each frame to one second, it allowed us to more reliably compare non-periodic and irregular behaviors that were ill-defined with the wavelet approach. Qualitatively similar results were obtained using 1680-ms intervals, indicating our results are robust to the choice of interval (Supplementary Figure 3C).

Both our approach and the wavelet approach define each point in time as a vector of all tracked coordinates in a temporal window centered at that point. If the same behavior is repeatedly performed at different points in time, in principle these time points should be near each other in this multi-dimensional space. To make this space more intuitive, the multi-dimensional space can be projected into two dimensions using t-Distributed Stochastic Neighbor Embedding (t-SNE)^5,6^. This projection requires a notion of movement similarity between pairs of 880-ms windows. We therefore defined a Posture Alignment Metric (PAM) that flexibly scores movement similarity in the presence of slight temporal and spatial variation in movement execution (Figure 3B), which was key to discovering stereotyped movements in the heterogeneous environment of the web. In this compressed space, frequently sampled behaviors form broadly consistent densities that can be segregated and partitioned in the reduced coordinate system^5,6^. Since anterior and posterior leg movements frequently act independently, we embedded anterior and posterior legs separately (Figure 3C). Anterior leg coordinates were oriented with respect to the midline of the prosoma, indirectly capturing three-dimensional body rotation, whereas posterior leg coordinates were oriented only with respect to the orientation of posterior legs in order to robustly pick up more subtle posterior movements (Supplementary Figure 3C). This approach produced consistent embeddings, with different spiders producing similar behavioral profiles in t-SNE space (Figures 3C,D, Supplementary Figure 3D).

To quantify the similarity of t-SNE embeddings, the Jensen-Shannon (J-S) divergence was used, since it does not rely on prior assumptions of density geometry^25^ (Figure 3D). When different webs were compared, the webs did not cluster based on the identity of the spider that produced the web (colored circles in Figure 3D), nor on the presence of stabilimenta or atypical web progression. The dendrogram branches from hierarchical clustering were fairly shallow, indicating the observed clusters were not necessarily well-separated. The outlier webs appeared to be due to excessive leg pausing (Supplementary Figure 3D), but the other web behaviors were fairly similar to each other based on the J-S metric, regardless of web geometry, typical/atypical web progression, individual, or stabilimentum occurrence.

To discriminate the t-SNE-defined behaviors more easily, a watershed function was applied to the t-SNE densities to partition them into discrete regions. Several of the behaviors in these regions could be easily annotated as distinct behaviors that had been observed previously, such as leg sweeps, abdomen bending, stabilimentum construction and silk-pulling, confirming the utility of this automated approach (Figure 3E, Supplementary Movie 2). Anterior legs performed a greater variety of exploratory-type movements. Since the spider builds the web blindly in complete darkness, frequent sweeps of the anterior legs were used to search for silk lines (Left, Right, and Alternating sweeps, Figure 3E). In addition, anterior legs participated in walking behaviors. Since anterior markers were influenced by the three-dimensional prosoma orientation, anterior limb trajectories also signaled rotation, abdominal bending, and stabilimentum behavior.

Posterior legs were primarily used for pulling and/or guiding silk from the abdomen. Even walking behaviors along the silk were primarily performed by the medial and anterior legs. Four general silk-pulling densities were observed, two of which corresponded to slow and fast silk-pulling (Figure3E: highlighted in yellow and green, respectively). The slower pulling was characteristic of the slower withdrawal and guiding of silk during proto-web, radii and auxiliary spiral stages, and occurred at a range of slow frequencies (Supplementary Figure 3D). Faster pulling was characteristic of the silk “combing” of capture silk observed previously for this genus^6^ and occurred at a fairly consistent leg frequency of 9 Hz (Supplementary Figure 3D). Subtle silk pulling characteristic of the stabilimentum stage could similarly be identified by posterior legs (pink highlight, Figure 3E). Occasionally silk-anchoring behavior could be identified by posterior legs, but posterior limb trajectories were more subtle during abdomen-bending compared to anterior limb trajectories due to the previously mentioned difference in their coordinate system (orange highlight, Figure 3E).

Due to the regular geometry of the web, several of these behaviors occurred at regular intervals across different webs. Anchoring, for example, regularly occurred when the spider anchored capture silk onto radii and therefore it frequently occurred at positions where the spider crossed radial lines (Figure 3F). Likewise, stabilimentum behaviors were detected almost exclusively where the spider built this structure (Figure 3F). The fast silk-combing is characteristic of capture silk construction^26^ and was exclusively observed at positions where capture silk was produced (Figure 3F). Alternatively, slower silk-pulling is used for non-capture silk, and thus only occurred at loci where non-capture silk was produced (Figure 3F).

The broad representation of identified behaviors across different webs, and the consistency with which they were performed during specific periods of web construction and at particular locations reinforces the accuracy of the automated method we employed to define web behaviors. Behaviors that have been manually characterized in the past such as capture thread combing, leg sweeps, and silk anchoring were easily discriminated with our method, which did not require manual annotation of movies. These behaviors were sampled at comparable levels across different web constructions, regardless of spider identity, web geometry, or typical/atypical web progression.

### Common and unique behaviors characterize each stage

Since the construction of different web features such as radii and capture-spiral are separated in both space and time (Figure 1F), behaviors that primarily occurred in certain web regions (Figure 3F) also showed strong temporal biases. This was more obvious when the behavior densities in t-SNE space were sub-divided by stages of web-building (Figure 4A-B). Anterior leg movements sampled during the proto-web and radii stage were very similar, as expected by the similar construction executed in both. Both stages were characterized by the construction of long lines spanning large parts of the arena, and thus more walking events occurred in these stages versus other stages (Figure 4B, blue highlight). Pauses occurred more frequently in these stages, as identified through either anterior or posterior limb dynamics (Figure 4B, gray highlighted regions). The proto-web in particular involved long pause periods (Figure 1I) that comprised ~90% of this web stage (Figure 1I). Radii construction also involved long pauses, though the durations were not as long (Figure 1I). The posterior densities were characterized by an absence of fast silk-combing which only occurs during capture spiral construction.

**Figure 4:**
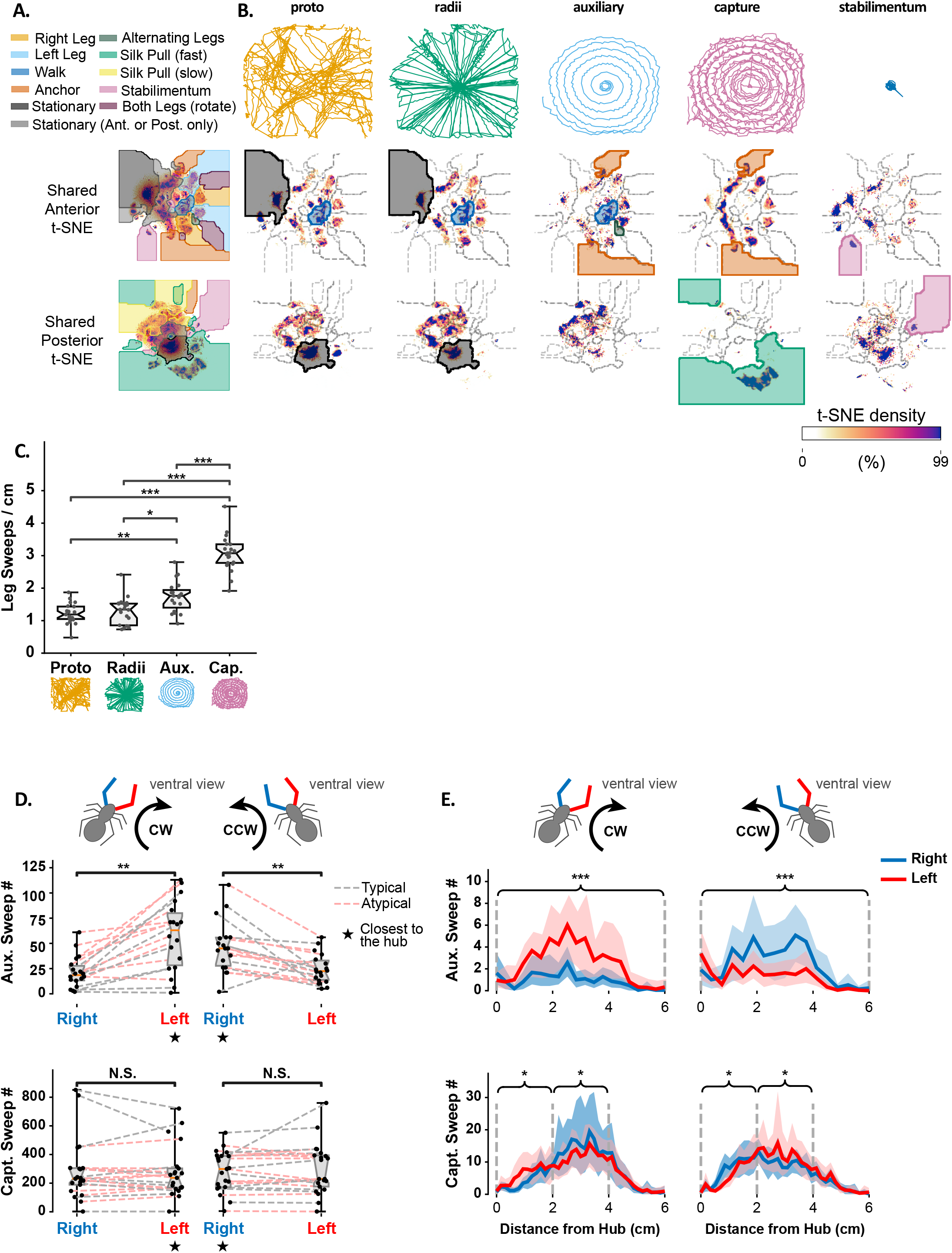
Stages of web-building use both shared and stage-specific motifs. A. Top: Legend of anterior and posterior movement motifs. Bottom: Cluster boundaries for each movement in the anterior and posterior embeddings. B. Top: Example centroid paths of a spider during different web stages. Middle: Anterior t-SNE embeddings for different web stages, averaged across all 21 recordings, and clipped at the 99th percentile to show structure. Shading used in A indicate motifs of note in the text. Bottom: Same as Middle, but for Posterior t-SNE embeddings. C. Spatial density of leg sweep occurrences by stage of web-construction. Notches represent the 95% confidence interval of the median. P-values based on Bonferroni-corrected two-sided Mann-Whitney rank tests (*: p < 0.05, **: p < 0.01, ***: p < 0.001). D. Left and right leg sweep occurrences during the auxiliary spiral (top) and capture spiral (bottom), for atypical (red) and typical (grey) recordings. CW: Clockwise, CCW: Counter-clockwise. Notches represent the 95% confidence interval of the median. P-values were computed using the Wilcoxon rank test (**: p < 0.01, N.S.: not significant). E. Left leg (red line) and right leg (blue line) sweep occurrences during the auxiliary spiral (top) and capture spiral (bottom) stages as a function of distance from the hub. Shaded regions represent the bootstrapped 99% confidence interval of the mean. P-values are computed using a Wilcoxon rank test by first pooling occurrences as indicated by the braces (*: p < 0.05, ***: p < 0.001). P-values for capture spiral comparisons were Holm–Bonferroni adjusted. Note that we repeated this analysis for un-pooled histogram bins with qualitatively similar results.

Auxiliary spiral construction behavior was similar to that of the proto-web and radii phases, but comprised significantly fewer pauses, and shorter pause duration (Figure 1I). While anterior walking movement occurred during this stage, rapid alternating leg sweeps were more common in this stage (olive green shaded region, Figure 4B). However, this stage saw an increase in anchoring behavior (dark orange region, Figure 4B) due to the fact the auxiliary spiral repeatedly crosses radii to which the auxiliary spiral is attached.

The increase in anchoring continued with the capture spiral construction, which also requires anchoring the capture spiral to radii. Like the auxiliary spiral, this stage also exhibited fewer and shorter pauses than the proto-web and radii stages (Figure 4B, 1I). Very little walking behavior was observed in this stage, as the spider is primarily focused on attaching capture spiral which is orthogonal to radii (Figure 4B, blue highlight). The anterior legs were often stationary (light gray region, Figure 4B), however these were not pauses per se, but the spider’s grasping radii while the posterior legs combed the silk. This combing behavior, which is characteristic of this stage^26^, was the most defining feature of this stage of web-building, as the posterior legs were nearly exclusively dedicated to this movement (green region, Figure 4B). If the spider built a stabilimentum, both the anterior and posterior legs moved in a rhythmic motion that was unique to this stage (pink regions, Figure 4B).

Leg sweeps were used at all stages of web building (light orange and light blue regions, Figure 4A) but occurred more densely in auxiliary spiral and capture spiral stages (Figure 4C). Since spiders build their webs blindly, almost exclusively through touch^1^, the increased leg sweeps correlate with a greater density of silk lines for the spider to touch and guide their movement. This is especially true during auxiliary and capture spiral construction, where prior observations have noted the frequent use of previously assembled spiral to guide the spider’s movement along the web^27^.

To investigate this behavior more closely, we analyzed leg sweeps separately during clockwise or counterclockwise spiral construction trajectories. *U. diversus* often changes rotational direction during spiral assembly, which enabled us to investigate leg-sweep bias in both directions for each spider. During auxiliary spiral construction, there was a clear bias for leg sweeps of the leg closest to the hub, regardless of angular trajectory (Figure 4D). During this phase, the spider spirals outward from the center, so the previously assembled auxiliary spiral is closer to the hub. The observed bias for hub-facing leg sweeps is likely due to the spider tapping this spiral to guide their movement. No such bias existed on average for the capture spiral (Figure 4D).

However, if we plotted leg-sweep bias as a function of distance from the hub, a dependence on hub-distance was observed for both spiral stages. The inward bias for leg sweeps during auxiliary spiral construction only appeared at a radial distance of 1-2 cm from the hub, likely reflecting the absence of an auxiliary spiral at shorter hub-distances. For capture spiral construction, an outer leg bias dominated the early part of capture spiral construction far from the hub, but then switched to an inner leg bias as the spider moved closer to the hub. This switch may reflect the spider initially using the prior capture spiral to guide assembly, but then switching to the inner legs where the radii silk density is higher to gauge distance from the hub.

The changing use of the legs during different phases of web building likely reflects the changing structure of the web, as well as the changing goals as the spider progresses through construction phases. Both auxiliary and capture spiral stages involve motion orthogonal to the radii, but the behaviors employed, and their temporal biases, were not necessarily the same. Besides the auxiliary spiral assembly occurring over a shorter timeframe (Figure 1F,H), this spiral construction also involved less stationary use of the anterior legs when compared to capture spiral construction. These results, in addition to the leg sweep biases, reflect changing coordination of behaviors during different web stages to create different structures in a similar context.

### Different behavioral transitions define stages of web building

Even though behaviors like leg sweeps were shared in different stages of web-building, how they were coordinated was not necessarily similar. Capture spiral construction in particular was the most highly regular. The posterior legs primarily performed silk-combing, followed by regular bouts of anchoring the capture silk to radii (Figure 5B, Supplementary Movie 4). The anterior legs either remained stationary or performed leg sweeps (Figure 5B). The regular progression of these behaviors can be seen when we plot the probability of their occurrences relative to the silk-anchoring behavior (Figure 5C). After anchoring the capture silk to radii, the silk-combing often starts slowly (yellow squares Figure 5C), followed by leg sweeps (orange and yellow squares, Figure 5C), brief pauses (black squares, Figure 5C), and then fast combing (green squares, Figure 5C). The regularity of these behavioral transitions reflects the regularity of the capture spiral, and the coordination that is needed to ensure proper construction of this web feature.

**Figure 5:**
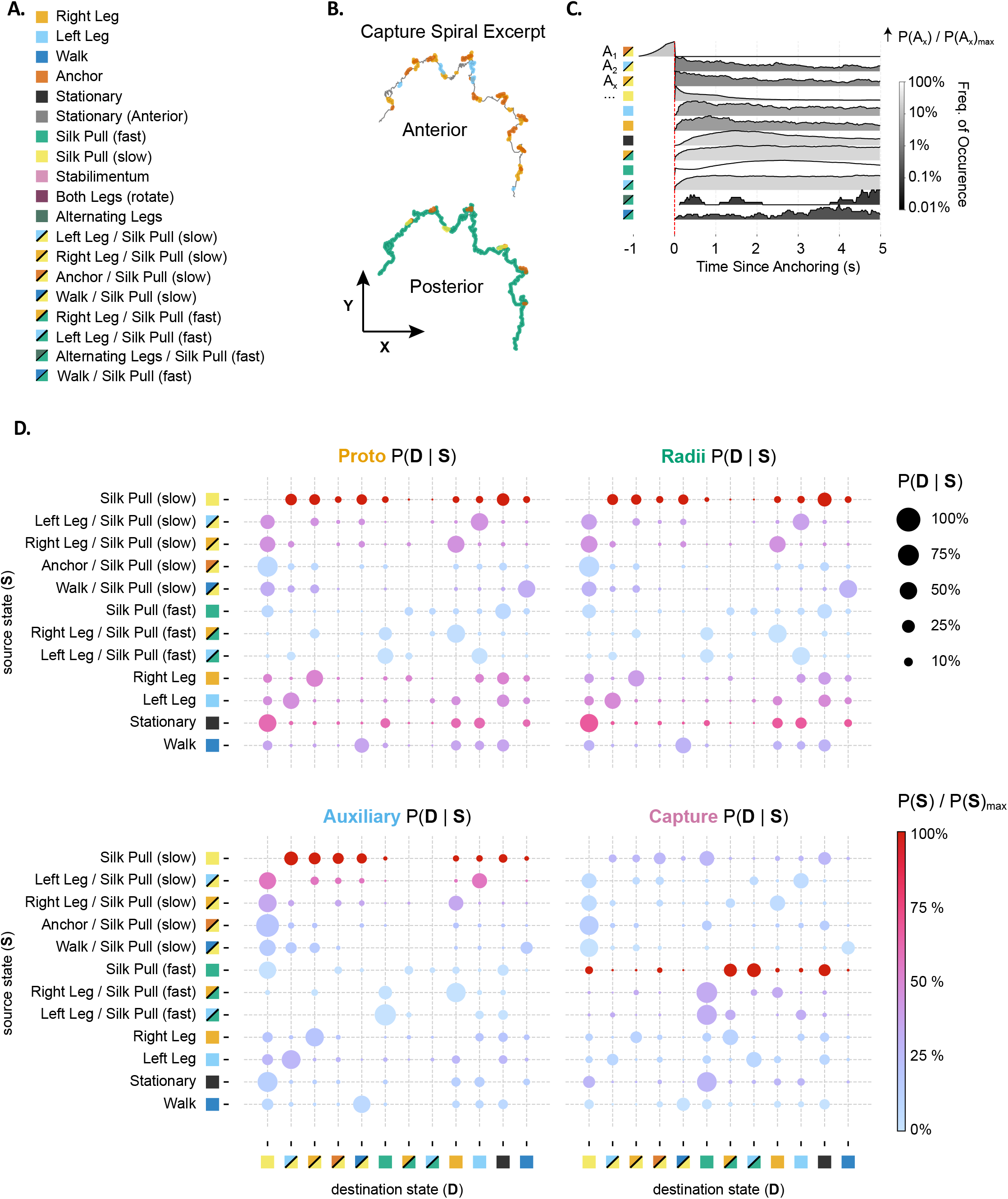
A. Legend of movement motifs in (B-D), representing a subset of movement motifs occurring across recordings. Single-color squares represent single-movement occurrences. Split-color squares represent combined anterior/posterior movements. B. Example trace of a spider’s trajectory during capture spiral construction. Each point is colored according to which anterior (left) or posterior (right) t-SNE density it accessed. C. Probability profile for movement motif timing relative to anchoring events during capture spiral construction. D. Transitional matrices for different web stages. Rows are starting behaviors (S), and columns are subsequent behaviors (D) immediately following behavior S.

Even though fast silk combing is unique to this phase, in principle other behaviors such as leg sweeps and anchoring should be coordinated differently during other web phases since the construction needs are different during each phase. When we constructed transition matrices for each web phase, many transitions were shared between stages and several were distinct. Since the spiders do not use their vision for web construction, posterior silk-pulling was often coupled with anterior leg sweeps (Figure 5D). These transitions from [leg-sweeps + silk-pulling] to [stationary anterior legs + silk-pulling] likely reflect the spider initially searching its local environment with its anterior legs, locating lines, and then holding on to them while the posterior legs continued pulling silk from the abdomen. This general transition was observed for both slow and fast silk pulling, and it was a behavioral rule which dominated the transition spaces for all phases. This is likely due to the need to navigate and continuously pull the silk from the abdomen during construction.

Certain transitions were more characteristic of certain stages and reflected more frequent occurrences of the source state. For example, transitions in and out of walking and turns were more likely in the proto-web and radii states, reflecting the frequent roles these behaviors played in assembling these structures. Likewise, fast extrusion transitions dominated the capture spiral stage. (Figure 4D).

Even though most behaviors are shared across stages, the differences in relative sampling of behaviors, and their transitional biases implies that each stage of web building is not only reflected in the spider’s trajectory, but in the behavior of the spider itself.

### Hierarchical Hidden Markov Model can predict web stages based on behavior-alone

If behavioral dynamics are truly distinct in different web stages, then we should be able to predict web stage by these behaviors alone. We investigated whether the changing transitional probabilities between behaviors could be used to categorize phases of web-building, independent of known spider position. We assembled a Hierarchical Hidden Markov Model (HHMM) with hidden top-level parent states and non-hidden child states representing the different movement motifs (Figure 6A, Supplementary Figure 6A). The parent states represented regimes with different underlying behavioral transitional probabilities, while the child-states represented the behavioral transitions themselves. No constraints were placed on state transitions within a regime, but transition from one movement state in regime X to one in regime Y could not occur without going through the parent transition state rXY, which reflects the probability of transitioning between regimes (Figure 6A, Supplementary Figure 6A). Models were repeatedly trained^28^ on a subset of data (in-sample) and used to predict regime changes in out-sample data (Figure 6B).

**Figure 6:**
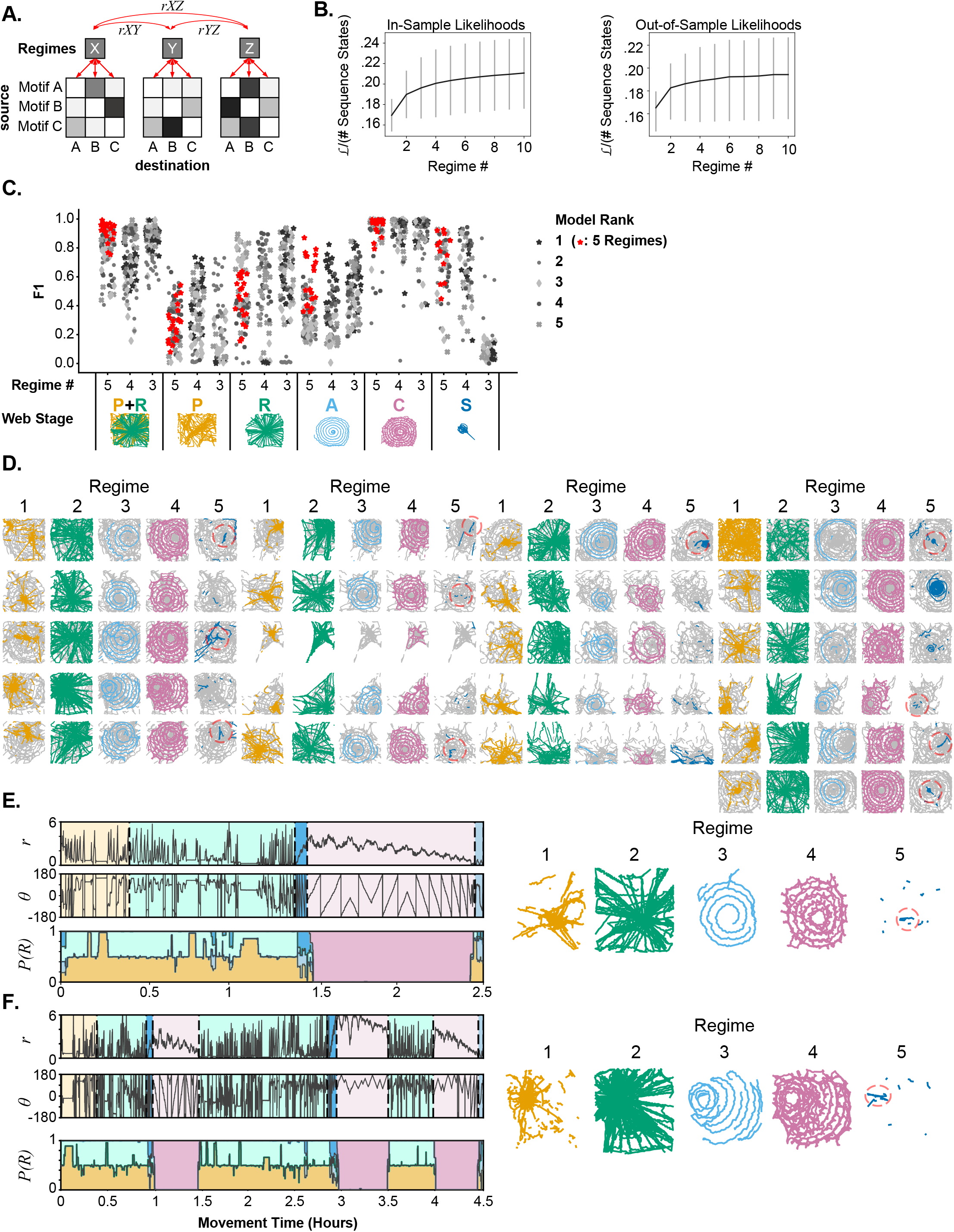
Stereotyped and distinct action sequences characterize stages of web-building. A. HHMM design. Within each parent state (regime), transitions are modeled as simple Markov processes characterized by a movement motif transition matrix (child states). Transitions between regimes (rXY, rXZ, etc.) by design can only occur through parent nodes, creating a hierarchical structure. B. Likelihood for in-sample (left) or out-sample (right) HHMMs for different regime numbers. Gray vertical lines represent the standard deviation of model likelihoods. C. F_1_ scores for the top 5 models for each regime-number limit. The top-ranked 5-regime model is highlighted in red, and used for D-F. Model accuracy (F_1_ score) defined as the agreement between manual stage annotations and stage boundaries predicted by each HHMM. F_1_ scores were defined based on the optimal mapping of HHMM predicted regimes to manual regimes. For models with fewer than 5 regimes, some predicted regimes were assigned to multiple web stages. The F_1_ score for the combined proto/radii stage was computed by first merging proto-web and radii annotations for both predicted and manual regimes. D. Spider positions for all regime occupations and webs for the 5-regime HHMM highlighted in red in (C). Spider positions were assigned to a regime based on the most likely regime designation for that time-point. Dashed red circles indicate stabilimenta. E. Web-stage manually predicted by spider position versus regime state (bottom panel) predicted by the 5-regime HHMM highlighted in red in (C) for a typical web progression. P(R) is the probability of each regime state. Centroid trajectories corresponding to the maximum-likelihood regime estimate are displayed (right panels), with colors matching the bottom panel. F. Same as (E), but for an atypical web progression.

As expected, the likelihood of the models increased with an increased number of regimes, however these likelihoods plateaued quickly after 3 regimes (Figure 6B). Due to the existence of local optima in this E-M fitting procedure, and the random initialization of each transition matrix before fitting, a variety of models were obtained. F1 scores^29^ were calculated for each model to assess how well these models matched manually annotated web stages based on the precision (true positives / true positives + false positives) and recall (true positives / true positives + false negatives) of the model.

When 5 regimes were chosen to match the number of web stages (Figure 1E), the 5 regimes accurately captured auxiliary and capture spirals for most webs, as well as the stabilimenta (Figure 6C). The discrimination between proto and radii stages was mixed, with regime 1 dominating periods with more pauses (Figure 6E). This is consistent with the underlying similarity of these phases (Figures 4B, 5D) and the difficulty in assigning a discrete transition between the phases^6^. Hence, the HHMM model also had difficulty discriminating between these phases, however as a combined state, these two regimes accurately captured proto-web/radii phases (Figures 6C-E). Prior to HHMM modeling, the stabilimentum stage was the least sampled behavior since it did not occur in all webs, and occurred over a short duration (Figures 1H, 4A). This stage mostly consisted of self-transitions (i.e. uninterrupted “stabilimentum” leg movements, Figure 4B), yet our HHMM model ignored self-transitions. Despite this, 5 regime models consistently identified stabilimenta stages in webs that had stabilimenta (Figures 6C-D). The categorization of most stages of web-building -- including proto-web/radii, auxiliary spiral, capture spiral, and stabilimentum -- occurred consistently across many recordings (Figure 6D), indicating that these stages are characterized by stereotyped behavioral transitions and constitute behavioral programs shared across individuals.

Regime transitions closely matched web-stage transitions defined by animal trajectory, even for animals that had atypical web-stage progression (Figure 6F). In the example given in Figure 6F, the spider started the typical progression of web building but then repeated a second web, which was again interrupted during capture spiral construction. Despite the atypical progression of web-building, the 5-regime HHMM accurately captured the proto/radii, auxiliary spiral, capture spiral, and stabilimentum stages. This indicates that the coordination of underlying behaviors was not necessarily altered, but instead that their spatiotemporal pattern of execution was atypical.

## Discussion

Orb webs are the result of fairly well-defined stages of construction that can be defined by a spider’s position. Unlike the static nature of the final web, tracking the spider’s trajectory allows us to monitor construction progress and compare differences in how spiders create webs that may appear similar in the end but are the results of different behavioral histories. Even though most webs are the result of a defined sequence of assembly stages, spiders can also perform an atypical progression through these stages. However, in the absence of more detailed knowledge of the spider’s behavior, the underlying behavioral differences that lead to atypical construction are difficult to uncover. Structural differences could be due to altered behaviors or shared behaviors which are performed in an atypical progression.

By tracking leg movements at a high spatiotemporal resolution, we were able to identify stereotyped anterior and posterior leg behaviors that were shared across spiders. Our choice of flexible alignment metric for movement comparison was key to detecting irregular but stereotyped movements across heterogeneous web environments. Whereas some movements such as abdomen bending were more homogeneous, others like slow silk pulling were more variable and structure-dependent, and future work may improve on the granularity of movement categorization. Though some behaviors, in particular posterior silk combing and manipulation during capture spiral and stabilimentum stages, occurred exclusively during those stages, most stages of web-building were not due to stage-specific behaviors, but rather a biased sampling of these behaviors. Furthermore, differences in spider trajectories, including those of atypical stage progressions, do not appear to result from differences in underlying behaviors. Leg sweeps were common in all stages as the spider blindly navigated the web, however how they were coordinated with other behaviors differed depending on web stage. Thus, different web-stages are the result of a mixture of unique and shared behaviors that are coordinated differently in each stage.

To explicitly quantify these differences in coordination, transition matrices for the probability of transitioning from one behavior to another were defined for each web stage. The transition matrices were unique for auxiliary and capture spiral, though the matrices for proto-web and radii were very similar. This was in keeping with the similarity of these behaviors when tracking the spider’s movement. Both stages involved the construction of long suspended lines, though the radii stage is the result of more regular radii construction. Also, by defining these stages in exclusive terms, we masked the gradual transition from proto-web to radii construction, since radii are assembled while the spider removes the proto-web. Despite performing a variety of similar actions in all stages of web-building, how spiders coordinate these actions differs depending on the construction stage.

Though the Hierarchical Hidden Markov Models, fitted to the behavioral sequences, were blind to the web-stages derived from the spider’s position and to the previously computed stage-specific transition matrices, they nonetheless mostly replicated these stage identifications, even for atypical web progressions. Interestingly, proto-web and radii stages were not categorized independently by our model, indicating these stages appear to sequence distinct movements in similar ways. This shows the difference in geometry likely arises from differences in where and when actions are executed —i.e., differences in sensorimotor rules — which our model and data currently does not capture. However, the successful identification of proto/radii, auxiliary spiral, capture spiral, and stabilimentum based on transitional probability alone shows that these resulting structures are physical records of underlying behavioral regimes.

The existence of regime-specific behavioral dynamics demonstrates that web-making is not a memoryless process; the sequence of behaviors a spider performs is dependent on the stage of web-building it occupies. Where this memory is stored, however, remains to be addressed. The web itself constitutes a physical memory of the behavioral state of the spider that together with purely reflexive sensorimotor rules could perhaps give rise to the observed phases of construction^30^. Alternatively, some or all of this memory could be stored as internal states in the brain. Prior work on black widow and funnel-web spiders has demonstrated the existence of path integration memory ^31, 32, 33^. In contrast to behavioral states such as satiety and arousal that are mostly hidden and need to be inferred indirectly, the physical outputs of the phases of web-building provide a useful record of the spider’s behavioral state, facilitating study of the external and internal factors generating these behavioral regimes.

Neuroactive chemicals such as caffeine and methamphetamine are known to alter certain parts of the web geometry, while leaving others intact^2^. These drugs alter neuromodulatory pathways in the brain, which indicates these pathways may participate in web-building behavior to varying degrees depending on web-stage. In addition to human manipulation, the larvae of several species of *Ephialtini*, a tribe of ectoparasitic wasps, can alter ecdysone levels in spiders to pause web-building prior to capture spiral construction, indicating neuromodulation may contribute to encoding the transition from radii to spiral constrution^34^. Similar internal states may regulate the progression of web phases even in the presence of ambiguous or noisy tactile sensory input, perhaps explaining the extraordinary robustness of this complex behavior in varying environmental contexts^35^.

Integration of sensory input and internal states is likely integral to the fidelity and robustness of web construction, though the relative contributions of reflexive and cognitive factors to this behavior remain to be addressed^10^. The simultaneous recording of both behavior and web structure would be needed to address the influence of local web architecture on behavior. Pharmaceutical delivery and subsequent behavioral analysis would be needed to infer how drug-induced changes in behavior lead to altered web architecture. The behavioral analyses performed here provide a useful set of tools to probe the underlying structure of this behavior further, and ultimately, how this structure is encoded in the brain.

---

**Supplementary Figure 1.A:**
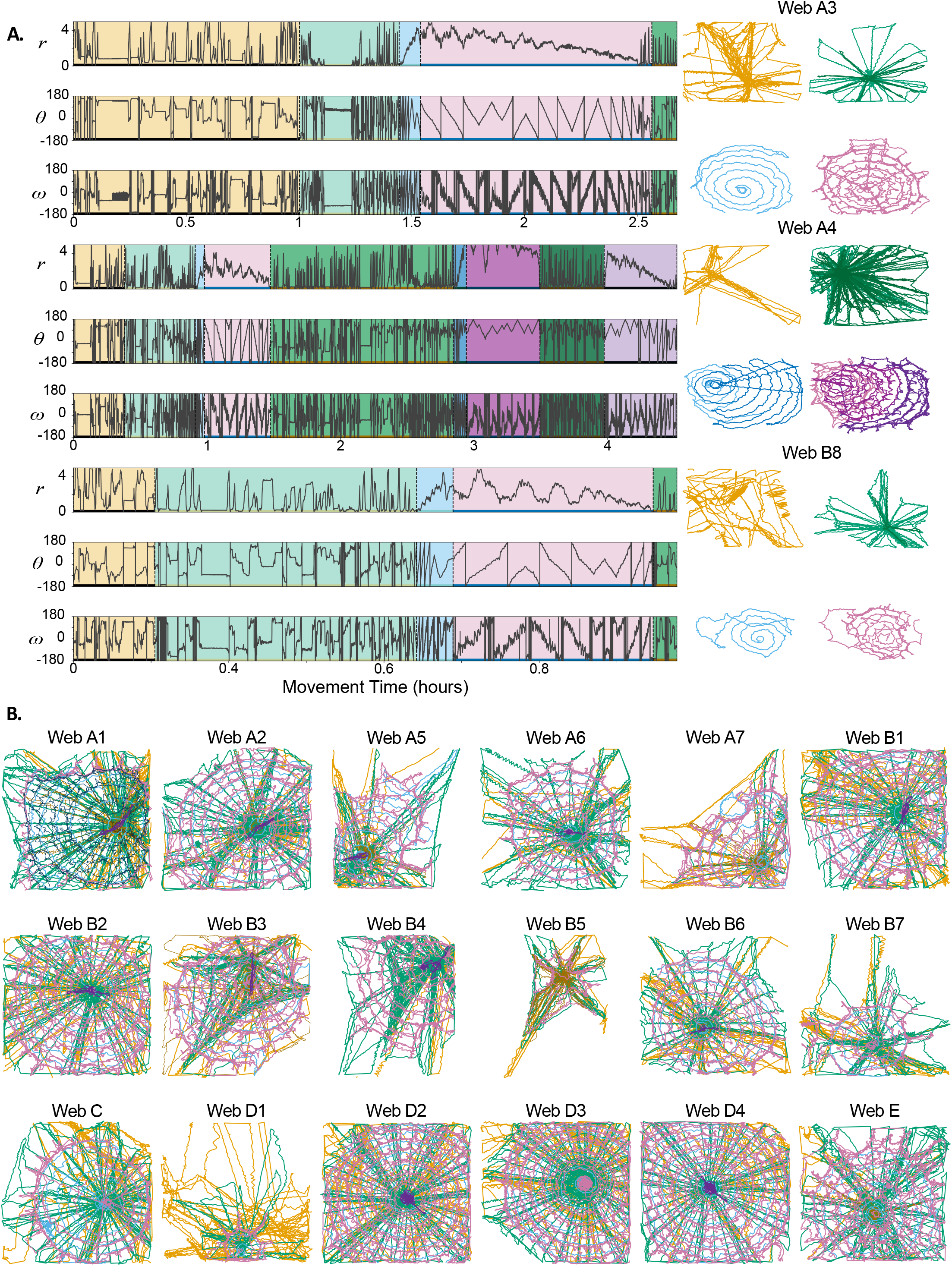
Behavioral diversity and similarity across all 21 recordings. A. Centroid trajectories for three additional recordings. B. Coordinates of web progression for the recordings not shown in (A). Webs are named by spider identity (Letters A-E) followed by the order in which each web was constructed over several days (Numbers).

**Supplementary Movie 1:** Movement trajectories from the web in Figure 1E-F.

**Supplementary Movie 2: Representative 1-second movement examples for each movement cluster**

**Supplementary Movie 3: Randomly sampled 1-second movement examples for each movement cluster, illustrating movement stereotypy and variation**

**Supplementary Figure 3.A:**
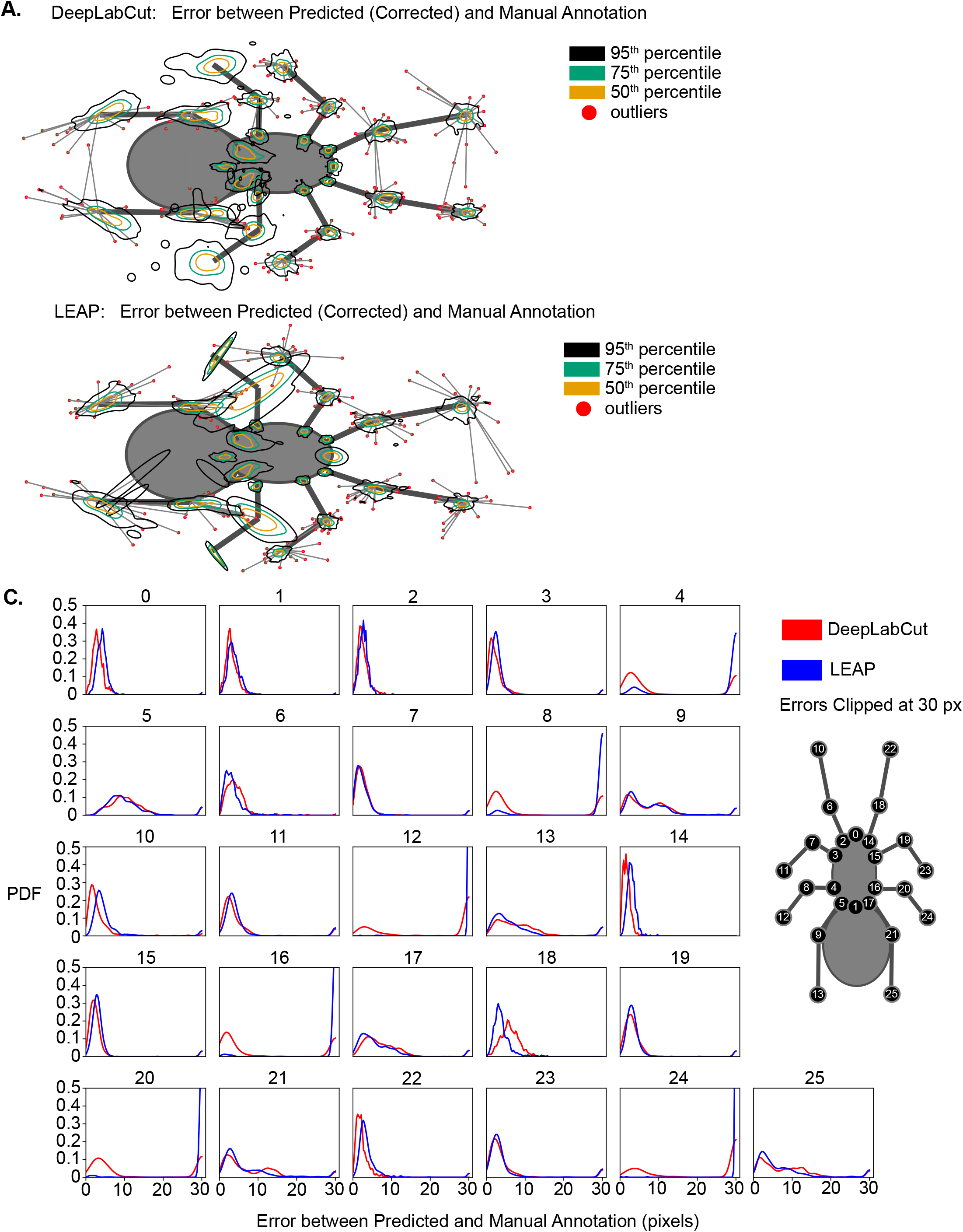
Overview of LEAP and DeepLabCut tracking performance. A. DeepLabCut confidence intervals, after limb tracking post-processing, reproduced from (3.A). Tracking error vectors are superimposed onto an arbitrary reference posture. 95th (black), 75th (green) and 50th (yellow) percentile contours are displayed. Errors outside the 95th percentile are displayed as red dots. B. LEAP confidence intervals, after limb tracking post-processing. Tracking error vectors are superimposed onto an arbitrary reference posture. 95th (black), 75th (green) and 50th (yellow) percentile contours are displayed. Errors outside the 95th percentile are displayed as red dots. C. Per-limb coordinate error histogram, before limb tracking post-processing, demonstrating the bimodal error distribution stemming partially from the occurrence of left/right limb confusion. Histograms are titled according to reference coordinates on the spider diagram to the right. PDF = Probability Density Function

**Supplementary Figure 3.B:**
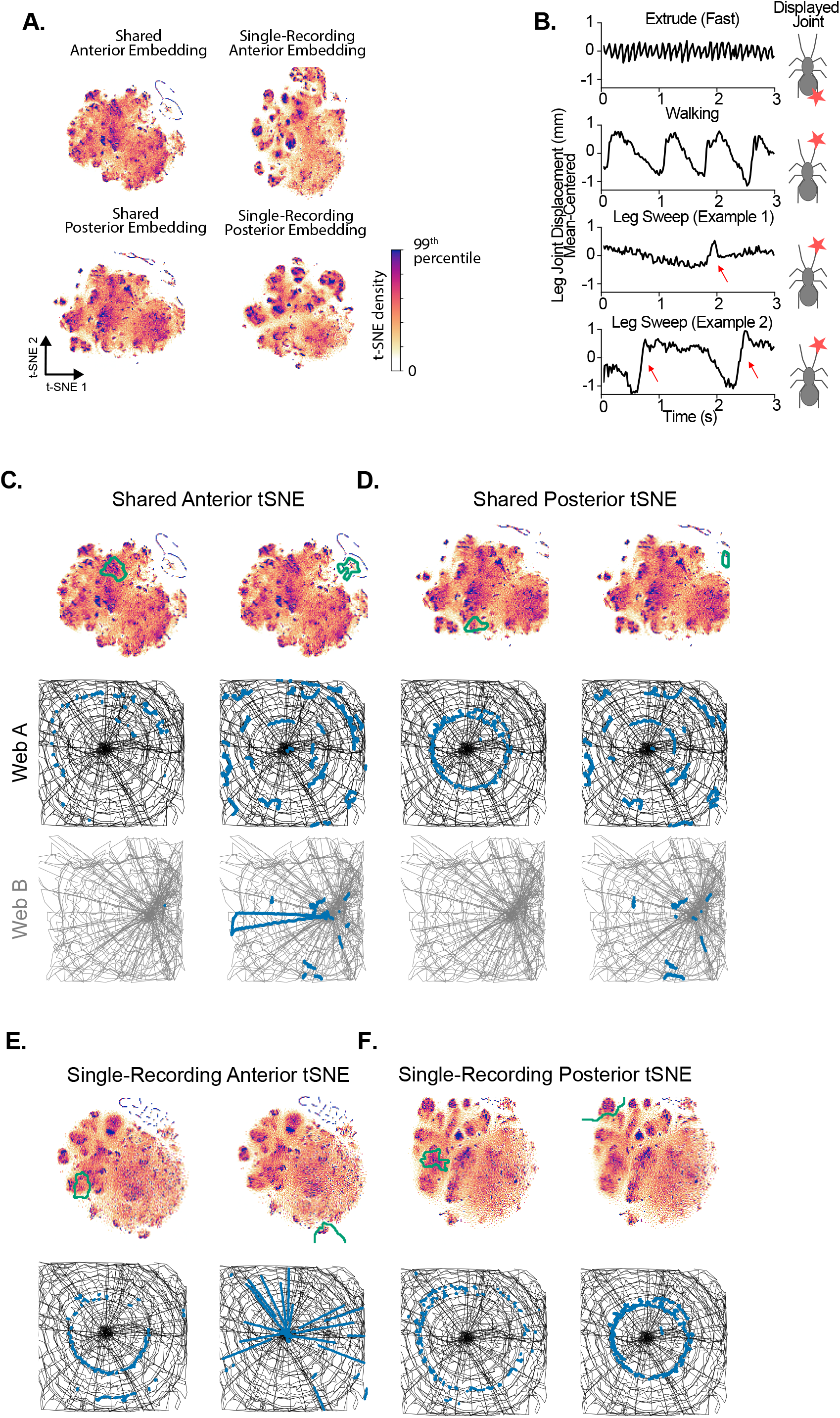
Movement clustering using wavelet embedding fails to identify shared movement motifs across recordings. A. Shared t-SNE embeddings (Left) versus an individual t-SNE embedding (Right). B. Examples of periodic movements (extrusion, top), occasionally-periodic movements (walking), and non-periodic movement (leg sweep, bottom). C. Top: Movement cluster outlines for two example anterior movement motifs. Bottom: Occurrences of behavioral motifs are highlighted in blue. The centroid history of two different spiders is in black and gray. D. Top: Movement cluster outlines for two example posterior movement motifs. Bottom: Occurrences of behavioral motifs are highlighted in blue. The centroid history of two different spiders is in black and gray. E. Same as C, but for t-SNE embedding obtained from a single recording. F. Same as D, but for t-SNE embedding obtained from a single recording.

**Supplementary Figure 3.C:**
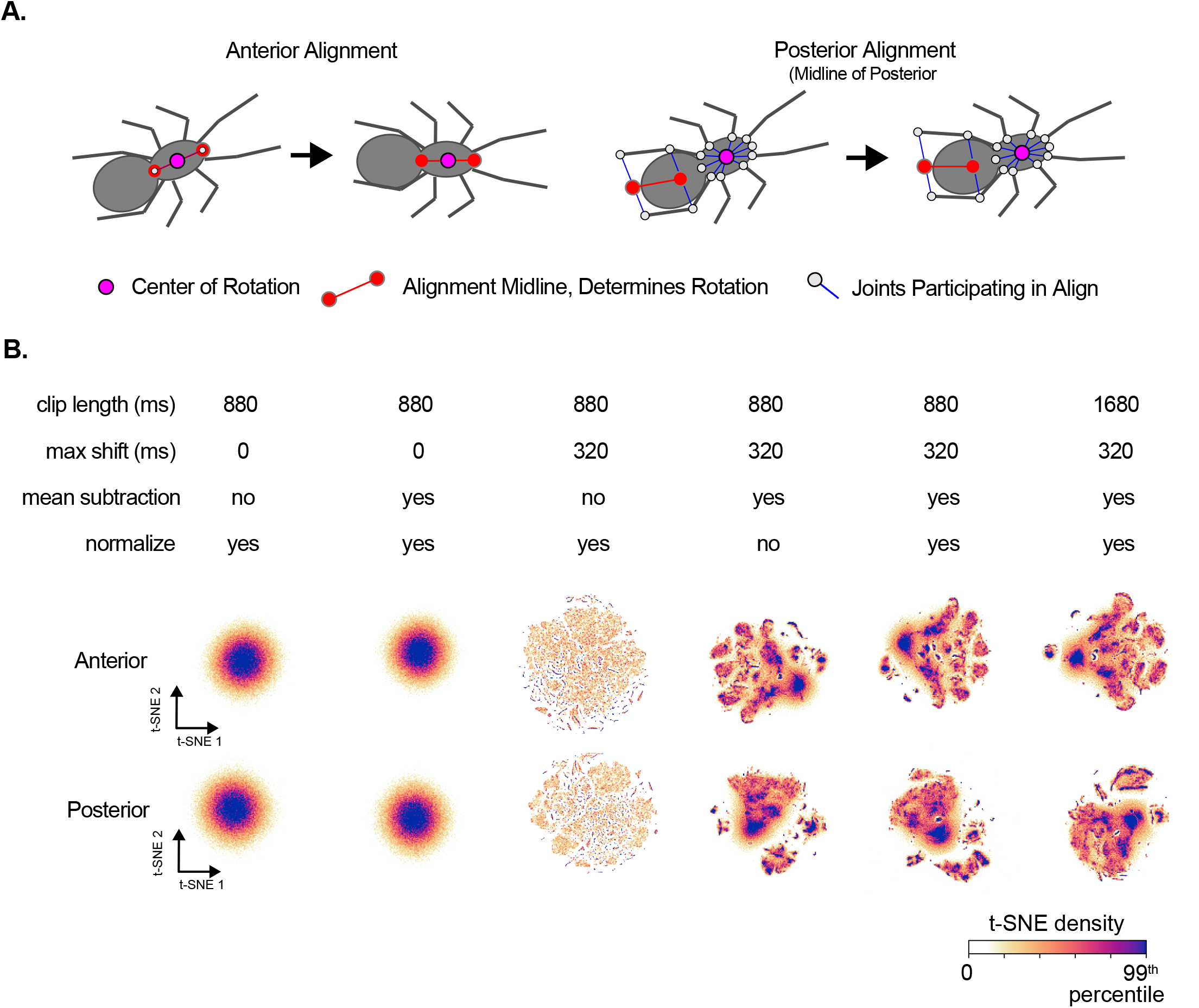
Posture Alignment Metric (PAM) methodology. A. Illustration of the body posture alignment used prior to applying the Posture Alignment Metric (PAM). Anterior leg joints are centered and rotated based on the anterior and posterior thoracic marker. Posterior leg joints are oriented with respect to the midline of posterior limb joints only. B. Overview of anterior and posterior shared t-SNE embeddings obtained with varying temporal and mean-shift parameters, indicating the importance of temporal realignment and mean-subtraction for movement clustering with PAM.

**Supplementary Figure 3.D:**
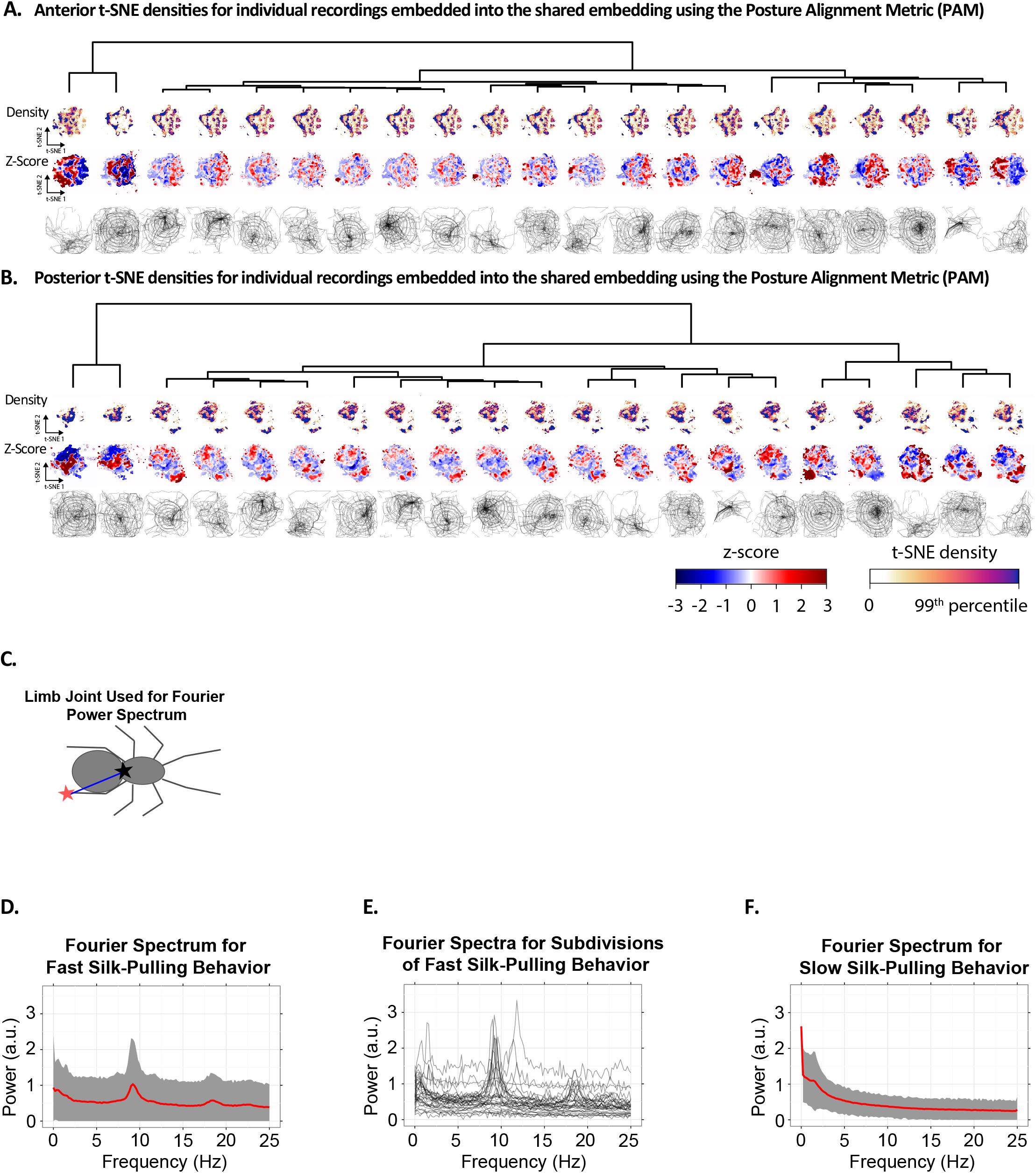
Hierarchical clustering of movement embeddings displayed in (Figure 3D) A. Hierarchical clustering of anterior movements using the Posture Alignment Metric (PAM). Each of the 21 recordings is displayed as a column, with its density (top), density Z-score with respect to the mean (middle) and web rendering (bottom). Densities are clipped to the 99th percentile before mean, standard deviation and Jensen-Shannon (J-S) distances are computed. Recordings are clustered based on pairwise J-S distances. B. Hierarchical clustering of posterior movements using the Posture Alignment Metric (PAM), as in (A). C. Limb Joint Diagram illustrating limb coordinate used in D-F. Fourier spectra are computed with respect to the displacement (blue line) of the left posterior limb joint (red star) relative to the posterior thoracic marker (black star). D. Fourier Power Spectra of the fast silk pulling behavior. Gray ribbons represent the mean ± standard deviation. E. Fourier Spectra for individual t-SNE subdivisions labelled as “fast silk-pulling” or “combing” behavior. Note the slight heterogeneity that exists in the amplitude and frequency of fast silk-pulling, explaining the subclusters that exist within the movements collectively labeled “fast silk-pulling” (Figure 3E). F. Fourier Power Spectra of the slow silk pulling behavior. Gray ribbons represent the mean ± standard deviation.

**Supplementary Movie 4:** Example capture spiral trajectory showing anterior and posterior t-SNE movement cluster timing.

**Supplementary Movie 5:** Example radii trajectory showing anterior and posterior t-SNE movement cluster timing.

**Supplementary Figure 6.A.:**
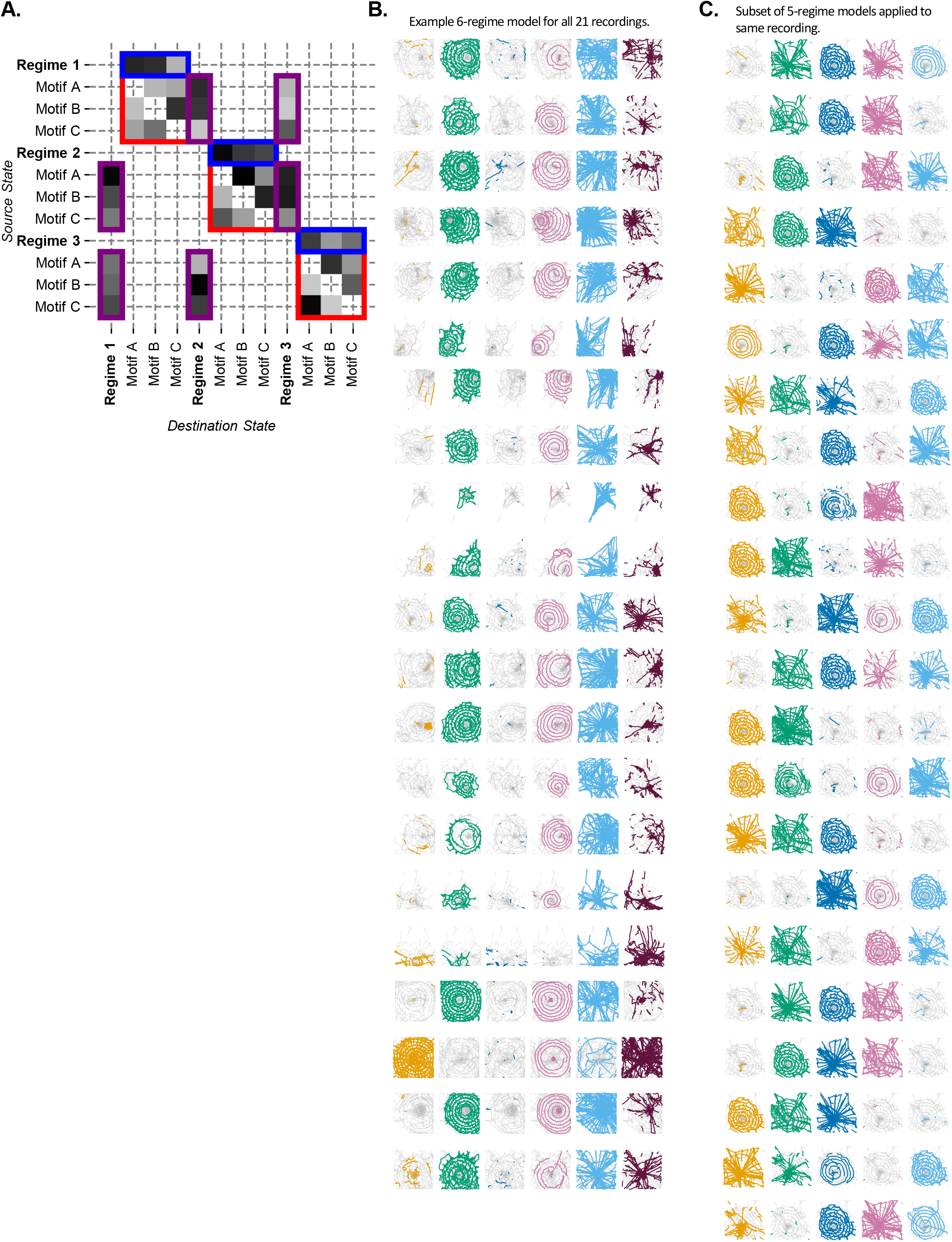
Hierarchical Hidden Markov Model (HHMM) structure. A. Example diagram of a Hierarchical Hidden Markov Model (HHMM) implementation as a constrained Hidden Markov Model (HMM). Red squares indicate within-regime transitions. Blue rectangles indicate transitions from hidden parent regime state (with no associated emission) to within-regime movement state. Purple squares indicate transition from a movement state to a parent regime state (with no associated emission). Note that a movement state cannot transition to its own parent regime state, to avoid loops. B. An example 6-regime HHMM, shown for all 21 recordings. C. A subset of 5-regime HHMM models for one recording. Note that only some model instantiations split out the auxiliary spiral stage. This is expected as the model fitting procedure encounters local optima, and models are initialized randomly. The large variation in model performance due to local optima is reflected in the large standard deviation in Figure 6B-C.

## Methods

### RESOURCE AVAILABILITY

#### Lead Contact

Further information and requests for resources should be directed to and will be fulfilled by the Lead Contact, Andrew Gordus.

#### Materials Availability

This study did not generate new unique reagents.

#### Data and Code Availability

All data were processed and analyzed using custom Python scripts, available at <www.github.com/GordusLab/Corver-Wilkerson-Miller-Gordus-2021>. The raw video recordings and data files supporting the current study have not been deposited in a public repository because of their large file size but are available from the corresponding author on request.

Our extension of the Flika^36^ GUI, which assists in visualization and exploration of the various datasets used in this study, is available upon request.

### EXPERIMENTAL MODEL AND SUBJECT DETAILS

Spiders of the species *Uloborus diversus* were collected in Half Moon Bay, CA. The population was housed in an on-campus greenhouse at Johns Hopkins University. All animals were transferred to custom indoor habitats and kept on a 12 hr:12 hr light-dark cycle (15-30 °C, 50-70% RH) at least two weeks before being used for behavioral experiments. Each habitat contained up to 9-12 individuals. Spiders were fed *Drosophila melanogaster* and *Drosophila melanogaster virilis* once a week. Only adult females were used in this study as adult males rarely build orb webs.

### METHOD DETAILS

#### Behavioral Assay

Adult females were recorded for an average of 24 hours in a custom behavioral recording rig. Recordings were stopped when the spiders had finished building the web. Spiders were transferred to a plexiglass arena with a 10 cm x 10 cm perimeter, coated with paper at the edges to encourage web-building. To increase the contrast of the spider with the background, we used a high-absorption background material below the behavioral arena (Acktar, Spectral Black Foil, SB-20×030-1-010). A camera (FUR FL3-U3-13Y3M-C, 1024×1024 pixel recording resolution) with a 35 mm fixed-focal length lens (Edmund Optics, #85-868) recorded their behavior at 50 Hz under 840 nm illumination by a ring light (Advanced Illumination, RL4260-880100LIC). The aperture of the lens was set to a minimum to maximize depth-of-field, and the fact that there was only one elevation at which the web could be built ensured the behavior was in focus throughout web-building. FLIR’s FlyCapture software was used to store the recordings in MJPEG 75% AVI format.

#### Manual Annotation of Web Stages and Hub Location

To manually segment the recordings into stages of web-building, as well as annotate the location of the hub, we built time lapse visualizations of the spider’s position history, which greatly facilitate discovery of the frame indices corresponding to boundaries between stages. Due to the large size of our recordings, storage constraints prohibited us from pre-rendering various visualizations. To facilitate exploration of our datasets, we therefore extended Flika^17^, an existing Graphical User Interface (GUI)^17^. Rather than pre-rendering our data visualizations, we have added GUI features allowing on-the-fly rendering of behavioral trajectories, allowing precise annotation of stage boundaries and hub location.

#### Limb Tracking

To facilitate improved limb tracking, each frame of the recording was first cropped and rotated to a 200×200 image to center and align the spider. The spider was detected heuristically, by thresholding the image and applying erosion/dilation cycles to remove the legs and find a body feature of expected size, after which the long axis of this feature was computed to determine the spider’s orientation. For most recordings, the image brightness was stereotyped enough that an arbitrary threshold of 30/255 could be used. For the recordings in which bright perimeter or background pixels were mistaken for the spider, 10,000 frames were randomly sampled and a pixel value histogram was computed at every x,y position. The lower peak of this histogram represents the mean and standard deviation of background brightness and was used to Z-score every frame prior to blob detection and alignment. In these cases, an arbitrary threshold of 20 standard deviations above the mean was used to detect the spider. As the raw AVI recording data was stored in batches of ~34,000 frames, these files were skipped if the spider was entirely stationary.

To analyze the movements of the spider, we tracked 26 points on its body: the base, femur and tibia of every leg, as well as the anterior- and posterior-most points of the prosoma. We randomly sampled 100,000 frames from one representative recording and used the Figure Eight AI service (Figure Eight Inc.) to obtain manual annotations. We then manually reviewed these annotations and selected 10,000 high-quality annotations for the next training step.

To automate limb annotation, we evaluated two CNN tracking frameworks: LEAP^21^ and DeepLabCut^22^. Both algorithms were trained on the 10,000 high-quality manual annotations. Both algorithms reached similar performance on the training sample (8.2 and 7.6 mean pixel error for LEAP and DeepLabCut respectively, 4.5% and 3.3% errors ≥25 pixels respectively). To evaluate out-of-sample prediction performance, we randomly sampled 456 frames across a total of 12 recordings, and created manual annotations. We excluded frames from manual annotation if the spider was touching the frame perimeter, or if the crop-rotate preprocessing step had incorrectly oriented the spider (rotated by 180 degrees). Both LEAP and DeepLabCut performed well, with a highly bimodal error distribution (Supplementary Figure 3A): limb joints were either highly accurately tracked, or often mistaken for a nearby limb joint, often on the opposite side. Because our tracking errors were driven mostly by this mismatch of limb joints, all subsequent analysis was done on the DeepLabCut tracking, due to its lower rate of large (≥25 pixels) errors (4.5% and 3.3% errors ≥25 pixels respectively for LEAP and DeepLabCut).

For subsequent analysis, periods without movement were excluded from the tracked dataset based on a conservative motion threshold (centroid velocity < 0.5mm/s). Due to camera parallax, the plexiglass arena created lower contrast of the spider on the background around the edges of the arena. Because this led to higher tracking errors, an iterative cropping procedure was used to shrink the effective frame size up to 50 pixels on each side of the frame until fewer than 10 above-threshold pixels existed at the edge of the cropped frame.

To minimize mis-tracked limbs, we detected mismatched joints by LEAP and DeepLabCut by monitoring limb lengths. When limb lengths increased beyond their representative lengths, indicating a tracking error, they were replaced by an interpolated position based on neighboring timepoints at most 200 ms away. If no recently available joint position was found, a linear regression model was used to impute a representative posture based on available joints. To detect erroneous limb lengths, we computed the mode (*Mo*) of the limb length distribution and computed the standard deviation (σ) based on only those limb lengths lying below the mode. Limb coordinates corresponding to limb lengths outside the range [*Mo-4σ, Mo+6σ*] were considered tracking errors and replaced as described.

#### Wavelet Transform

We applied the Morlet continuous wavelet transform to capture limb dynamics, as previously described^23^. Before the wavelet transform was applied, all limb coordinates were Z-scored. The spider’s posture was oriented with respect to the anterior and posterior thoracic markers (Supplementary Figure 3C). The frequency range used was 1 - 25 Hz, with 20 frequencies spaced as follows:

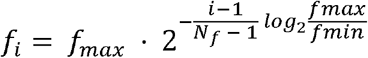

The wavelet transform was defined as follows, with ω_0_ = 5, as described by Berman et. al.^23^:

#### Posture Alignment Metric (PAM)

We sought an alternative to the wavelet transform as a means of discovering stereotyped movement motifs for more irregular, less periodic behaviors. The Posture Alignment Metric (PAM) directly captures the similarity between two movement motifs by attempting to optimally align the limb joint trajectories of each movement clip. Movement clips with fixed body orientation were sampled from each recording using a rolling window (880 ms, 44 frames). To capture the similarity of movement dynamics, we allow for slight differences in starting posture by mean-centering each movement clip. After mean-centering, limb coordinates are divided by the overall standard deviation of the limb coordinate across all frames. To ensure overall movement similarity is not obscured by slight differences in limb movement onset, limb coordinate time series are allowed to shift by up to 320 ms (16 frames) to minimize alignment error. Alignment error is computed as the summed Euclidean distance between all pairs of limb coordinate trajectories. Because the temporal realignment procedure is not symmetric, the alignment is repeated twice: once temporally shifting the first clip and holding the second one fixed, and once vice-versa. The resulting Euclidean errors are averaged, ensuring the PAM distance metric is symmetric.

Even though this procedure predetermines the timescale of the behaviors of interest, we experimented with different window lengths. There were no qualitative differences in our results when using a window length of 1680 ms (84 frames) instead of 880 ms, indicating this procedure is robust to slight variations in window length.

#### Embedding and Stereotyped Movement Identification

A successful categorization of movement motifs considers not only the relative similarity of movement occurrences, but also their frequency of occurrence. To decompose web-making behavior into a smaller set of stereotyped movement motifs, we used a dimensionality reduction approach to discover frequently sampled movement patterns^5^. Since the anterior and posterior legs contain the majority of movement information, and are most robustly tracked, we limited our analysis to anterior and posterior leg movement. When both anterior and posterior movements were embedded together, anterior leg movements often dominated the alignment metric and caused mixed posterior movements to be clustered together. To improve the discrimination of posterior movements, anterior and posterior leg movements were embedded independently.

To compare limb movement trajectories independent of the absolute rotation of the spider, the anterior leg coordinates were aligned using the midline of the thorax (Supplementary Figure 3C). Note that even though we recorded limb trajectories in two dimensions, due to three-dimensional body rotation, this thorax-alignment procedure captured body rotation resulting from, for example, abdomen bending during silk attachment.

Although this thorax midline alignment helped capture three-dimensional behaviors, it complicated the separation of anterior and posterior movements, as both reflect three-dimensional body rotation. Because this complicates the detection of subtle stereotyped posterior leg movements, we instead aligned posterior leg movements based on the midline of posterior limb coordinates. More specifically, for a given posterior limb joint, a midpoint was computed as the average of the left and right instance of this joint. Each frame was then reoriented to minimize the maximum distance of any mid-point to the x-axis (Supplementary Figure 3A). This isolated the orientation of posterior leg joints from three-dimensional body rotation and anterior leg movement, thus improving the separation of anterior and posterior movement motifs.

In order to discover stereotyped movements across recordings and individual spiders, we randomly sampled 200,000 timepoints collectively from all recordings. We used the OpenTSNE^37^ (version 0.3.11) Python package to embed these timepoints into two dimensions. For wavelet transform embeddings, the wavelet transform was first applied to the Z-scored limb trajectories, as described above. We used the Euclidean distance as the distance metric. For Posture Alignment Metric (PAM) embeddings, movement clips were extracted using a rolling window, as described above. The PAM was used directly as the embedding metric.

We repeated embeddings with a variety of parameters, including a range of perplexities (50 – 500), but found the resulting embedding to be quite robust to the choice of parameters. In our subsequent analysis we used a perplexity of 100. We also experimented with other dimensionality reduction methods including UMAP^38^ (version 0.3.8) but found the results to be qualitatively similar for the purposes of movement clustering, and used t-SNE throughout our subsequent analyses.

We next looked for frequently sampled stereotyped movements in the form of local maxima in this dimensionally-reduced space. We computed a two-dimensional histogram of the 200,000 embedded timepoints and smoothed this histogram by convolving each data point with a Gaussian proportional to the average distance to its 10 nearest neighbors in t-SNE space. Local maxima were computed for the resulting smoothed histogram and watershed was performed to segment the t-SNE space into stereotyped movement clusters, as previously described^5^.

Having identified shared movement motifs across recordings, the embedding procedure was repeated for each individual recording, using the previously computed shared t-SNE embedding space, and the previously obtained shared watershed segmentation was used to determine the movement cluster for all individual recording frames.

To facilitate our understanding of the embedding space and clustered movements, we further extended the previously mentioned Flika GUI to display t-SNE embeddings side-by-side with movement clips and web renderings.

#### Manual Simplification of Movement Motifs

The unsupervised movement embedding and classification generated 91 anterior and 85 posterior movement clusters. This movement classification is too fine grained for our subsequent analysis, as it would result in excessive computational complexity of subsequent modeling steps, and generally complicates intuitive understanding of behavioral transitions. We therefore manually reviewed and grouped the previously generated movement motif clusters. We randomly sampled movie clips from each recording centered on the occurrence of a given movement cluster. After manually reviewing the clips, we identified a subset of anterior — *left leg sweep, right leg sweep, both legs (rotate), alternating legs, walk, stabilimentum, anchor, all legs stationary, anterior legs stationary* — and posterior movements — *stabilimentum, anchor, fast silk-pulling, slow silk-pulling, all legs stationary, posterior legs stationary*. If a cluster produced predominantly mixed, unstereotyped movements, it was left unlabeled. 84% of anterior clusters received labels, and 92% of posterior clusters received labels, or similarly, 87±4.7% of frames received anterior labels, and 94±2.5%% of frames received posterior labels. Manual labeling was blind to the position of the cluster in t-SNE space, yet almost all manual labels formed contiguous areas in t-SNE space, indicating that clusters close in t-SNE space indeed represent highly similar movement motifs.

To facilitate the transition matrix analysis and HHMM analysis, we converted the separate anterior/posterior movement cluster identities into a single, merged movement state. The merged movement state was defined simply as the combination of the anterior and posterior movement state, with a few exceptions: If either the anterior or posterior embedding signaled a whole-body *stationary* state, the merged state was reduced to *stationary*. If the anterior movement state was unlabeled or indicated the anterior legs only were stationary, only the posterior movement state was used — and vice versa. Finally, only anterior legs were used to signal the stabilimentum state, as this movement is most consistently detected by the anterior leg embedding, due to the anterior legs being aligned using the thoracic midline which captures body orientation.

#### Hierarchical Hidden Markov Model

To explore the existence of behavioral regimes characterized by different transitions between movement motifs, we trained a Hierarchical Hidden Markov Model (HHMM) on our data. In our model, only the transition regime state is hidden. Within a given transition matrix regime, the behavior is modeled as a Markov process without hidden states, with each manually annotated movement motif corresponding to a given Markov state.

In order to exclude noisy transitions, a movement motif was only included if the spider spent at least 240 ms in that (previously manually simplified) cluster. To speed up training, we simplified the model by removing self-transitions from all datasets. This reduced the entire web-making behavior to an average of 12,322 transitions (7,427-15,854 transitions 10-90th percentile).

To fit the model, a Hierarchical Hidden Markov Model (HMM) was constructed using the Python library Pomegranate^28^ (version 0.11.1). To model the Markov process corresponding to a given regime, a regime-specific “hidden state” was created for each movement motif which deterministically emitted the corresponding movement motif. Although any movement-to-movement transition was allowed within a given regime, transitions from a movement motif in one regime to a movement motif in another were only allowed to occur by going through a non-emitting parent state (Supplementary Figure 6A). These transition matrix constraints therefore effectively generate the hierarchical nature of the model.

For model training, the recordings were divided into 5 groups, and all models were trained using 5-fold cross validation. Every model was trained at least 50 times (10 times per fold) with random initialization, and trained using the Baum-Welch algorithm. Due to computational resource constraints, we trained every model for at least 250 iterations, with improvement ratios falling to less than 1%.

To quantify the agreement between the manual web-building stage boundaries and those determined by the HHMM, we computed precision, recall and F_1_ statistics for each HHMM model and recording. The predicted regime was defined as the regime with the highest probability at a given timepoint. To capture the agreement of long-duration states, we used the rolling mode of the predicted state over a 30 second window. Periods of long pauses, defined as centroid trajectories that remained within a 1 mm radius for 30 seconds, were not considered for the F_1_ statistic. Because the mapping of HHMM-predicted regimes to manual annotation regimes is arbitrary, we chose the mapping for each HHMM model that maximized the F_1_ statistic. For HHMM models with fewer than 5 predicted regimes, some predicted regimes were assigned to multiple manually annotated regimes. Statistics for the combined proto-web/radii stage were computed by first merging both the manual and predicted proto-web/radii stages for the given model. To highlight the 5 best-performing models for a given number of predicted regimes, we defined overall model performance as follows: For each model, corresponding predicted-to-manual regime mapping, and recording, we computed the worst-case F_1_ performance across the proto-web/radii, auxiliary spiral, capture spiral and stabilimentum. For the 3-regime models, only the proto-web/radii, auxiliary spiral and capture spiral were used to assess performance. The median performance across recordings was then computed for each model and corresponding predicted-to-manual regime mapping. We subsequently selected the predicted-to-manual regime mapping for each model that maximized its median F_1_ score. This yielded one F_1_ score for each of the 50 fitted models for each regime count, of which 5 are displayed.

### QUANTIFICATION AND STATISTICAL ANALYSIS

All analysis was performed using custom Python 3.6.7 scripts. All statistics reported as X±Y represent mean±standard deviation unless otherwise noted.

Cumulative travel distance was computed based on the movement of the center of the thorax in 50 frame (1 second) increments, with only movement steps above 0.5mm/s counted towards this distance metric. Travel distances by stage were computed based on manual annotations of stage boundaries. Total stage durations were similarly computed based on manual annotation of stage boundaries, except for the start of the proto-web, which was defined as the point at which 10% of the total proto-web distance was traveled. This more accurately reflects true construction duration of the stage, as the start of web-building is strongly circadian-dependent.

#### Pause Definitions

The quantification of pause duration by stage was computed using two strategies. First, if a range of frames was detected as being stationary based on centroid velocity (<0.5mm/s), a t-SNE embedding cluster was not computed. Second, if a range of frames passed this cutoff, but the computed t-SNE embedding for either the anterior or posterior legs signaled a stationary posture, this frame range was marked as a pause. All other frames were marked as non-pause states. Pauses for a given stage were computed over the [10%, 90%] interval of distance traveled for that stage. Pauses between stages were defined as those pauses occurring after 90% of the length of the prior stage and before 10% of the length of the subsequent stage was traversed.

#### Leg-Sweep Characterization

The quantification of leg sweep frequency by stage were computed as follows. Leg sweep occurrences were counted when the embedding was stable in a leg-sweep state (*left leg, right leg*, or *alternating legs)* for at least 240 ms. Contiguous movement durations beyond 240 ms were not re-counted until another non-leg sweep movement occurred. To more specifically count bouts of leg probing, we only counted occurrences at least 1 second apart, although the results are qualitatively similar without this constraint. To account for differences in web-size, we normalized this figure by the distance traveled in the given recording and stage of web-building. Web stage distance was computed as the sum of distances between the spider’s position every 1 second (50 frames), but distance segments were only summed if they were above an arbitrary noise floor (5 pixels/1 s) and below a maximum speed (500 pixel/1 s) that filters occasional tracking errors. Left-versus-right leg sweep frequencies show a dependence on the distance of the spider to the hub. To quantify this distance-dependent leg sweep bias, we manually selected candidate intervals for comparison and compared total leg sweep occurrences within these intervals based on Wilcoxon rank tests. In an alternative analysis, we determined leg sweep histogram bins based on the Freedman-Diaconis rule and tested for evidence of leg sweep bias within each bin using a two-sided permutation test. After Bonferroni-correction, this again revealed a leg sweep bias for a number of bins overlapping with our manually selected distance-from-hub intervals.

#### Radii Analysis

For all radii analyses, radii were determined as follows. We incrementally searched for centroid trajectory segments within the manually annotated radii stage that covered a minimal radial distance of 7 mm unidirectionally towards or away from the hub and passed within 2 cm of the manual annotation of the hub location. If a radius candidate segment came within 2 mm of the hub, it was split into two segments. Radii segments were further selected to have a maximum deviation less than 1mm from the straight line fit through that segment. The straight-line fit error was computed as the maximum deviation of the trajectory segment to the straight-line approximation. The angular position of radii was defined by the best-fit line of the radial segments through the hub, again defining the error of fit using the maximum deviation of the trajectory from the straight line. For our analysis of consecutive radii pairs, we selected pairs of consecutive outward (away from the hub) and inward (towards the hub) radii segments. To capture the gradual change in radii pair angular span over the course of web-building, we computed a rolling minimum and maximum angular span between radii pairs over a rolling window of 5 radii pairs. In the computation of neighboring radii spacing, to prevent double-counting re-traversals of the same radial lines, we merged previously detected radii segments using an agglomerative clustering approach with complete linkage and a ±7 degree cutoff.

## Acknowledgements

We thank D. Gordus for assistance in identifying field-sampling sites for *U. diversus*, K. Branson for advice in experimental design, M. Filipovitz for manual limb tracking, C. Bargmann, A. Bendesky, W. Eberhard, S. Flavell, P. Kidd, A. Sordillo, D. Ventimiglia, members of the Johns Hopkins University Biology and Neuroscience Departments and Gordus lab members for helpful discussions and comments on the manuscript. J.M. acknowledges funding from the NSF Graduate Research Fellowship Program. A.G. acknowledges funding from NIH (R35GM124883).

## Author contributions

A.C., N.W., J.M., and A.G. designed research. A.C. performed centroid-tracking analysis, post-limb-tracking analysis, and tracking fidelity analysis. N.W., J.M., and A.G. assembled the behavioral arenas. N.W. performed behavior experiments, and N.W. and A.C. performed limb-tracking and limb-tracking fidelity analysis. A.C. wrote all final software used in this manuscript. A.C. and A.G. analyzed the data and wrote the paper.

